# IRI-CCE: A Platform for Evaluating CRISPR-based Base Editing Tools and Its Components

**DOI:** 10.1101/2021.07.25.453636

**Authors:** Rahul Mahadev Shelake, Dibyajyoti Pramanik, Jae-Yean Kim

## Abstract

Rapid assessment of CRISPR/Cas (Clustered Regularly Interspaced Short Palindromic Repeats/CRISPR associated Cas protein)-based genome editing (GE) tools and their components are critical aspects for successful applications in different organisms. In many bacteria, double-stranded breaks (DSBs) generated by CRISPR/Cas tool generally cause cell death due to the lack of an efficient non-homologous end-joining pathway and restricts its use. CRISPR-based DSB-free base editors (BEs) have been applied for precise nucleotide editing in bacteria, which does not need to make DSBs. However, optimization of newer BE tools in bacteria is challenging owing to the toxic effects of BE reagents expressed using strong promoters. Improved variants of two main BEs capable of converting C-to-T (CBE) and A-to-G (ABE) have been recently developed but yet to be tested in bacteria. Here, we report a platform for *in vivo* rapid investigation of CRISPR-BE components in *Escherichia coli* (IRI-CCE) comprising different combinations of promoters/terminators. We demonstrate the use of IRI-CEE to characterize different variants of CBEs (PmCDA1, evoCDA1, APOBEC3A) and ABEs (ABE8e, ABE9e), exhibiting that each independent BE has its specific editing pattern for a given target site and promoter type. Additionally, the IRI-CCE platform offers a rapid way to screen functional gRNAs. In summary, CRISPR-BE components expressed by promoters of different strengths in the IRI-CCE allow an analysis of various BE tools, including cloned biopart modules and gRNAs.

## 1 Introduction

The adoption of the bacterial immune system, CRISPR/Cas (Clustered Regularly Interspaced Short Palindromic Repeats/CRISPR-associated protein), for targeted genetic manipulation has transformed the field of genome editing (GE) research (Anzalone et al., 2020). The CRISPR-based tools allow direct or indirect programmable engineering of all three major building blocks of life, i.e., DNA, RNA, and protein (Pramanik et al., 2021a). Fusion of deaminase with Cas enzyme forms a powerful GE tool called base editor (BE) that allows precise nucleotide conversion in the targeted region. The BE tools are based on the use of nickase (nCas9) or dead Cas9 (dCas9) (mainly from *Streptococcus pyogenes*) that does not need to make double-stranded breaks (DSBs) in the DNA for nucleotide substitution (Komor et al., 2016; Nishida et al., 2016). In animals and plants, primarily nCas9-BE is employed to stimulate the nicking of non-deaminated strands that produces single-stranded breaks (SSBs) and favor desired BE outcomes by tweaking the cellular DNA damage responses (Huang et al., 2021). The single guide RNA (sgRNA) together with the nCas9-BE or dCas9-BE complex binds the protospacer adjacent motif (PAM) in the target locus to form an R-loop, in which the partially open non-target strand (NTS) provides a possible substrate for base conversion. Depending on the type of deaminase used, editing can occur within or near the gRNA region, described as deamination or editing window (Rees and Liu, 2018).

In eukaryotes, DSBs generated by CRISPR/Cas tool are repaired by error-prone non-homologous end-joining (NHEJ) pathway leading to the gene disruption. Most prokaryotes lack the efficient NHEJ pathway, thus restricting the DSB-dependent CRISPR/Cas applications (Tong et al., 2021). The DSB-free BE tools are one of the best choices for site-directed precise base editing studies in prokaryotes. However, toxic effects of BE components under the control of strong promoters were earlier reported in bacteria, including *Escherichia coli* (Banno et al., 2018; Rodrigues et al., 2021). Using the protein degradation (LVA) tag together with dCas9 facilitated the BE tests in *E. coli*. In this regard, promoters expressing the optimal amount of BE components and sgRNAs may help to avoid cell toxicity and investigation of BE tools. Some virus-derived plant promoters have been reported to drive the transcription of downstream coding genes in *E. coli* (Jacob et al., 2002; Jopcik et al., 2013). For example, the transcriptional machinery of *E. coli* can recognize Cauliflower Mosaic Virus (CaMV) 35S promoter (p35S) (Assaad and Signer, 1990; Lewin et al., 1998), which is also the most often used promoter for expression of *Cas* gene in plant GE. Although the p35S is less active, this aspect is of great significance for rapid validation of cloned plasmids in *E. coli* or to design a heterologous expression platform consisting of differential-strength promoters. The use of validated CRISPR-BE biopart modules using Golden Gate assembly protocols (Engler et al., 2014) might be helpful for both prokaryotic and eukaryotic base editing experiments.

Here, we compared different promoters for their transcriptional activity in *E. coli* (**Figure 1**). Subsequently, a BE system was employed to establish an efficient and rapid evaluation platform to assess the differential-strength promoter-driven base conversion in *E. coli*, termed *in vivo* rapid investigation of <u>C</u>RISPR-BE <u>c</u>omponents in *<u>E</u>. coli* (IRI-CCE). We report that the efficient sgRNA expression was driven by the consensus sequence of Arabidopsis U6 promoter (pAtU6) (75 bp) in *E. coli* cells. The modular cloning approach described in the present work enables the validation of designed bioparts for further modular cloning of CRISPR-BE plasmids and the interchangeable use of bioparts independent of their context in the IRI-CCE platform.

**Figure 1.**
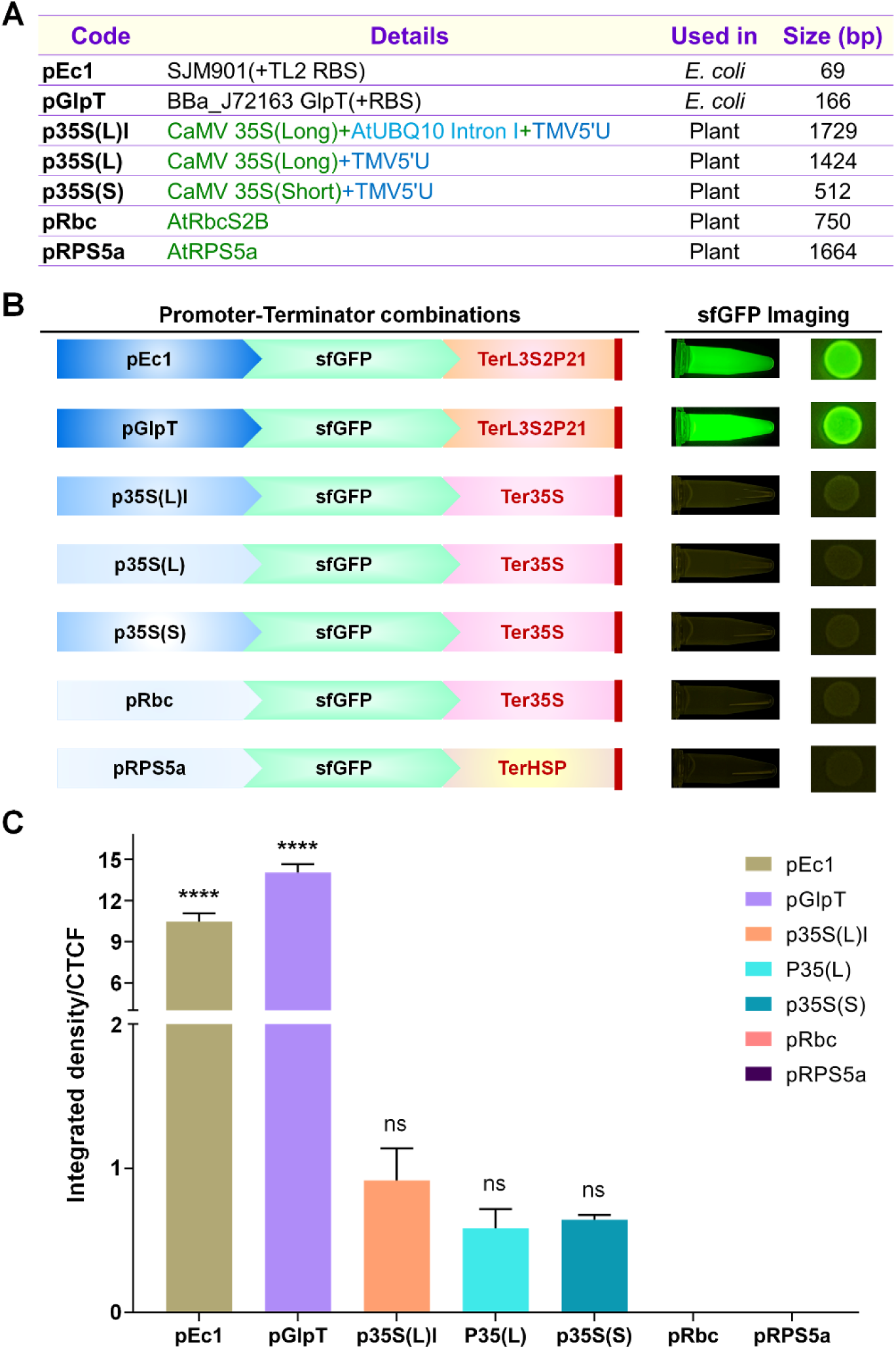
Screening of different promoters to evaluate the protein expression activities. **(A)** Information of bacterial, virus-derived plant, and native-plant promoters, along with their additional components and size used in the present study, is summarized. The details about the source are provided in Supplementary Information. (**B)** Schematic representation of superfold GFP (sfGFP)-expressing transcriptional units and sfGFP fluorescence in 10-beta *E. coli* cells. (**C)** Relative expression levels of sfGFP driven by seven different promoters were inferred from the corrected total cell fluorescence (CTCF). Statistical analysis of three independent biological replicates was performed and plotted using GraphPad Prism 9.0 software. The significant differences were calculated by one-way analysis of variance followed by Tukey’s multiple comparisons test and are indicated at the top (****P<0.0001, ns- not significant). Bars represent mean, and error bars represent the standard error of the mean (SEM) values.

Different versions of cytidine base editor (CBE) and adenine base editor (ABE) were analyzed for their substitution efficiency from C-to-T and A-to-G, respectively. Comparison between the differential-strength promoter-driven editing activities revealed the differences in the length of editing windows of BEs, including CBEs [Target-AID system based on sea lamprey cytidine deaminase, PmCDA1 (Nishida et al., 2016); evolved variant of PmCDA1, evoCDA1 (Thuronyi et al., 2019); human APOBEC3A, A3A (Zong et al., 2018)] and ABEs [ABE8e (Richter et al., 2020); ABE9e (Yan et al., 2021)]. Also, we demonstrate the potential use of the IRI-CCE platform for screening functional gRNAs by comparing the sets of active and inactive gRNAs. Moreover, editing outcomes of all the analyzed gRNAs imply that the promoter choice is a crucial factor in achieving improved BE or variable length of editing windows.

## 2 Methods

### 2.1 *E. coli* strains

The four *E. coli* strains were used: 10-beta, DH5α, DB3.1, and BL21(DE3). The information about strain genotype provided in **Supplementary Table 1**. The primers and plasmids used in this study are listed in **Supplementary Tables 2 and 3**, respectively. The *E. coli* strains were cultured in Luria-Bertani (LB) broth with appropriate antibiotics.

### 2.2 Plasmid construction and cloning

The required bioparts were amplified by conventional polymerase chain reaction (PCR) using a Phusion high-fidelity DNA polymerase (Thermo Fisher Scientific, Waltham, MA). The cloning of different plasmid vectors was designed and performed by following the principle of MoClo (Weber et al., 2011) and Golden Gate assembly (Engler et al., 2014) protocols using BpiI/BsaI Type IIS enzyme digestion-ligation. The DNA recognition sites of restriction enzymes BsaI and BpiI were removed from internal sequences of bioparts during PCR amplification to make them suitable for Golden Gate cloning (a procedure also termed domestication). The DNA oligonucleotide pairs for the pEc1 (SJM901+RBSTL2) promoter and TerL3S2P21 terminator were annealed and ligated into BpiI-digested acceptor plasmids. Various promoters including pGlpT, p35S(Long), p35S(Short), pRbc, and pRPS5a were used from the pYTK001 (Addgene #65108) (Lee et al., 2015), pICH51266 (Addgene #50267) (Engler et al., 2014), pICH51277 (Addgene #50268) (Engler et al., 2014), pICH45195 (Addgene #50275) (Engler et al., 2014), and pKI1.1R (Addgene #85808) (Tsutsui and Higashiyama, 2017), respectively. The promoter DNA sequences and their evaluation provided in **Supplementary Figures 1, 2, 3 and 4**. Similarly, DNA sequences of Terminators, Ter35S, and TerHSP were adopted from the pICH41414 (Addgene #50337) (Engler et al., 2014) and pKI1.1R (Addgene #85808), respectively.

Three cytidine deaminases: PmCDA1 (Target-AID) (Nishida et al., 2016), evoCDA1 (Thuronyi et al., 2019), and APOBEC3A (Zong et al., 2018), were PCR amplified from PmCDA1-1x uracil-DNA glycosylase inhibitor protein (UGI) (Addgene #79620), evoCDA1 pBT277 (Addgene #122608), and A3A-PBE-ΔUGI (Addgene #119770), respectively. For all three deaminases, the same linker regions were used as reported in the previous studies. The ABE8e was synthesized from Bioneer Co. (Daejon, Korea), and the ABE9e variant (containing additional V82S/Q154R mutations) was cloned using ABE9e module as a template by site-directed mutagenesis PCR. PmCDA1-1xUGI was fused to the C terminus of nCas9. The evoCDA1, A3A, and ABE8e were fused to the N terminus of nCas9 with the XTEN linker. The 2xUGI module (template: Addgene #122608) was fused to the C terminus of evoCDA1-nCas9 and A3A-nCas9. The sgRNA expression was driven by bacterial pJ23119 (synthetically cloned) or plant AtU6 promoter (pICSL01009, Addgene #46968). The nCas9(D10A) was generated by the PCR method using previously optimized Cas9 as a template (Level 1 hCas9 module, Addgene #49771). Desired sgRNA sequence was PCR amplified using plasmid pICH86966::AtU6p::sgRNA_PDS (Addgene #46966) as a template and cloned together with either pJ23119 or pAtU6 for sgRNA expression.

For cloning of target region of the desired sgRNA sequence, a pair of oligonucleotide DNAs that contains the target sequence with PAM was annealed and ligated into BsmBI-digested universal target-acceptor plasmid L1 or L2 having optimized superfolder green fluorescent protein (sfGFP) (Lee et al., 2015) at downstream side (**Supplementary Figure 5 and 6**).

### 2.3 Bacterial transformation, plasmid isolation, and Sanger sequencing

Ligation products of all the steps during the cloning were transformed into competent cells of *E. coli* 10-beta strain by a heat-shock method. The bacterial culture was spread on the LB media containing desired antibiotic and incubated at 37°C for 18-24 h. The cloned plasmids and BE activities were confirmed by Sanger sequencing at Solgent Ltd. (Daejeon, Korea) or Cosmogentech Ltd. (Seoul, Korea). For mutagenesis assay, the different *E. coli* strains were transformed with the appropriate plasmids using the heat-shock method and were pre-cultured for 1 h with 1 ml of LB medium. After incubation for 1 h at 37°C, the fraction of cell cultures were spread on LB agar (1.5%) plates containing selection antibiotics with needed concentrations. The next day, individual colonies from the plate were inoculated in 3 ml LB broth with appropriate antibiotics and cultured at 37°C. The culture time was varied according to the experimental parameters and mentioned in appropriate sections. The plasmid isolation was done using Plasmid Mini-Prep Kit from BioFact Co. Ltd. (Daejeon, Korea) for Sanger sequencing analysis.

### 2.4 Promoter activity analysis

As shown in **Figure 1B**, all the tested promoters were cloned with sfGFP sequence at the downstream side, followed by a termination signal. Bacterial cultures were grown from the transformed single colonies in LB broth, shaking at 37°C for 24 h. The grown cultures were incubated for 24 h at 4°C, and then OD_600_ values were normalized to 1. The cells were harvested at 10000 rpm for 1 min and washed with 1x PBS (Phosphate buffered saline). Images were captured in blue light with a FluoroBox from CELLGENTEK Co., Ltd. (Deajeon, South Korea). The fluorescence intensity level was quantified using the ImageJ software (Schneider et al., 2012), and corrected total cell fluorescence (CTCF) was calculated using the following formula: CTCF = Integrated Density - (Area of selected cell x Mean fluorescence of background readings).

For qRT-PCR, RNA was extracted using the following method. *E. coli*-carrying plasmids (**Figure 1B**) were grown in LB medium. Cells were harvested during the exponential growth phase (12 h with OD_600_ value 0.5). Total RNA was extracted using RNeasy Protect Bacteria Mini Kit from Qiagen. For all samples, 600 ng of total RNA was used for complementary DNA (cDNA) synthesis using a QuantiTect Reverse Transcription Kit from Qiagen following the manufacturer’s instructions. To estimate the relative *sfGFP* transcript, the quantitative real-time PCR (qRT-PCR) reactions were carried out using the KAPA SYBR FAST qPCR kit from Kapa Biosystems (MA, USA) with *sfGFP*-specific primer sets (**Supplementary Table 4**). Cycling of PCR consisted of pre-denaturation at 95°C for 5 min followed by 40 cycles of a denaturation step at 95°C for 10 min, an annealing step at 60°C for 15 s, and final extension step at 72 °C for 20 s using the CFX384 Real-Time System from Bio-Rad, Hercules (CA, USA). The qRT-PCR reactions were performed with independent biological replicates. Relative *sfGFP* transcript values normalized against internal control 16S ribosomal RNA (*rrsA*) gene. Data analyses were performed by the 2^−ΔΔCt^ method (Livak and Schmittgen, 2001).

### 2.5 Evaluation of editing activities

The single colonies were cultured, and the plasmid vectors containing synthetic targets were purified using Plasmid Mini-Prep Kit from BioFact Co. Ltd. (Daejeon, Korea) for further sequencing analysis. For sequencing analysis of genomic loci, the target fragments were PCR amplified using target-specific primers from the randomly picked colonies and then analyzed by Sanger sequencing. Sanger sequencing data analysis was performed using SnapGene software (GSL Biotech; available at snapgene.com). The editing efficiency was determined either by the ratio of non-edited to edited colonies from the randomly picked cells or by the nucleotide conversion rate using the online tool EditR (Kluesner et al., 2018) from C-to-T and A-to-G for CBE and ABE, respectively. The data were statistically analyzed and plotted in GraphPad Prism 9.0.0 (www.graphpad.com).

## 3 Results

### 3.1 Different promoters drive the gene expression with differential strength in *E. coli*

BE activity differs according to the expression level of BE components; therefore, we sought to test a set of bacterial, virus-derived plant, and native-plant promoters (**Figure 1A, Supplementary Figure 1**) for transcriptional activities in *E. coli*. We constructed sfGFP-expressing transcriptional units using an appropriate combination of promoters and terminators (**Figure 1B**). The transformed 10-beta *E. coli* cells were grown and amounts of sfGFP fluorescence were plotted in **Figure 1C**. The bacterial promoter pGlpT showed the highest expression levels among all the tested promoters. No activity was observed in the case of two native-plant promoters (pRbc and pRPS5a). All three p35S-based promoters expressed a comparable amount of sfGFP, consistent with the previous reports (Assaad and Signer, 1990; Lewin et al., 1998). Based on the level of sfGFP fluorescence, the promoter order from highest to lowest strength was determined as pGlpT>pEc1>p35S(L)I>p35S(S)>p35S(L)>pRbc/pRPS5a.

Further, relative mRNA transcripts of *sfGFP* were measured using qRT-PCR that showed a consistent pattern to that of protein amount except for pEc1 (**Supplementary Figure 2**). Interestingly, pEc1 showed maximum transcriptional activity among all the tested promoters. However, the mRNA abundance of *sfGFP* was not correlating with the fluorescence activity of sfGFP protein for pEc1, possibly due to poor mRNA-protein correlations at the analyzed time-point (Liu et al., 2016). pGlpT and p35S(L)I showed significantly higher activities than p35S(L), pRbc, and pRPS5a. Overall, pGlpT and p35S(L)I promoters exhibited the highest protein expression levels among the tested promoter groups, respectively.

### 3.2 Assessing PmCDA1-mediated C-to-T conversion driven by different promoters in *E. coli*

Apart from the selectable markers, the CRISPR-based BE construct consists of two major transcriptional units: deaminase-fused Cas9 mutant and sgRNA expression unit, driven by two separate promoters. Previous bacterial BE studies consisted of a deaminase fusion with partially impaired nCas9 or inactive dCas9 (Banno et al., 2018; Zheng et al., 2018; Zhang et al., 2020; Rodrigues et al., 2021). We chose the nCas9(D10A)-based CBE system in the present study because of the frequent use in BE studies in different organisms. Partially active nCas9 generates the SSBs that favor desired BE outcomes (Huang et al., 2021). To avoid the lethal effect of nCas9-BE, we opted to explore different promoters for the expression of CRISPR-BE machinery in *E. coli*.

We first sought to inspect ten different RNA polymerase III (RNA Pol III)-dependent promoters for conserved -10 (TATAAT) and -35 (TTGACA) elements (**Supplementary Figure 3**). The promoters were chosen because of earlier studies reporting efficient sgRNA expression in different species, including bacteria (pJ23119), plants (pAtU6, pOsU3, pOsU6, pMtU6.6, pZmU3, pTaU3, pTaU6), yeast (pSNR52), and human (phU6). The pJ23119 promoter has successfully been used for sgRNA expression in *E. coli* and other bacteria (Qi et al., 2013; Banno et al., 2018; Rodrigues et al., 2021). We detected partial or fully conserved -10 and -35 elements in all the promoters. Remarkably, the DNA sequence alignment of pJ23119 and pAtU6 showed 54.28% identity with fully conserved -10 and partially conserved -35 elements (**Supplementary Figure 4**). Therefore, in view of developing a heterologous promoter for sgRNA expression, pAtU6 was examined for optimal sgRNA expression in *E. coli*.

We employed a PmCDA1-based Target-AID system that induces C-to-T mutations in the editing window located approximately 1 to 8 bases from the distal end of the PAM (counting positions 21-23 for PAM) in 20 bp gRNA (Nishida et al., 2016). We adopted two approaches for BE testing. Firstly, a two-component system consisted of a target plasmid and a CRISPR plasmid including Pro-sgRNA and Pro-nCas9-PmCDA1-Linker-1xUGI (hereafter nCas9-PmCDA1) (**Supplementary Figure 6A**). Secondly, a single-component system composed of the target, sgRNA, and nCas9-PmCDA1 assembled into a single plasmid (**Supplementary Figure 6B**). We analyzed both single- and two-component plasmid vector systems, showing similar editing outcomes. Therefore, a single-component system was used for further analysis (**Figure 2A**). Next, two gRNAs were tested containing alternate Cs at even (Test gRNA1, **Figure 2B**) and odd (Test gRNA2, **Figure 2C**) positions spanning from 1 to 10 in the 20 bp gRNA. The target regions for the Test gRNAs were synthesized and cloned into the designed universal target-acceptor module (**Supplementary Figure 5**). Cloning of the required target region into a universal target-acceptor using MoClo kit protocol (Engler et al., 2014) permits the assessment of any intended gRNA.

**Figure 2.**
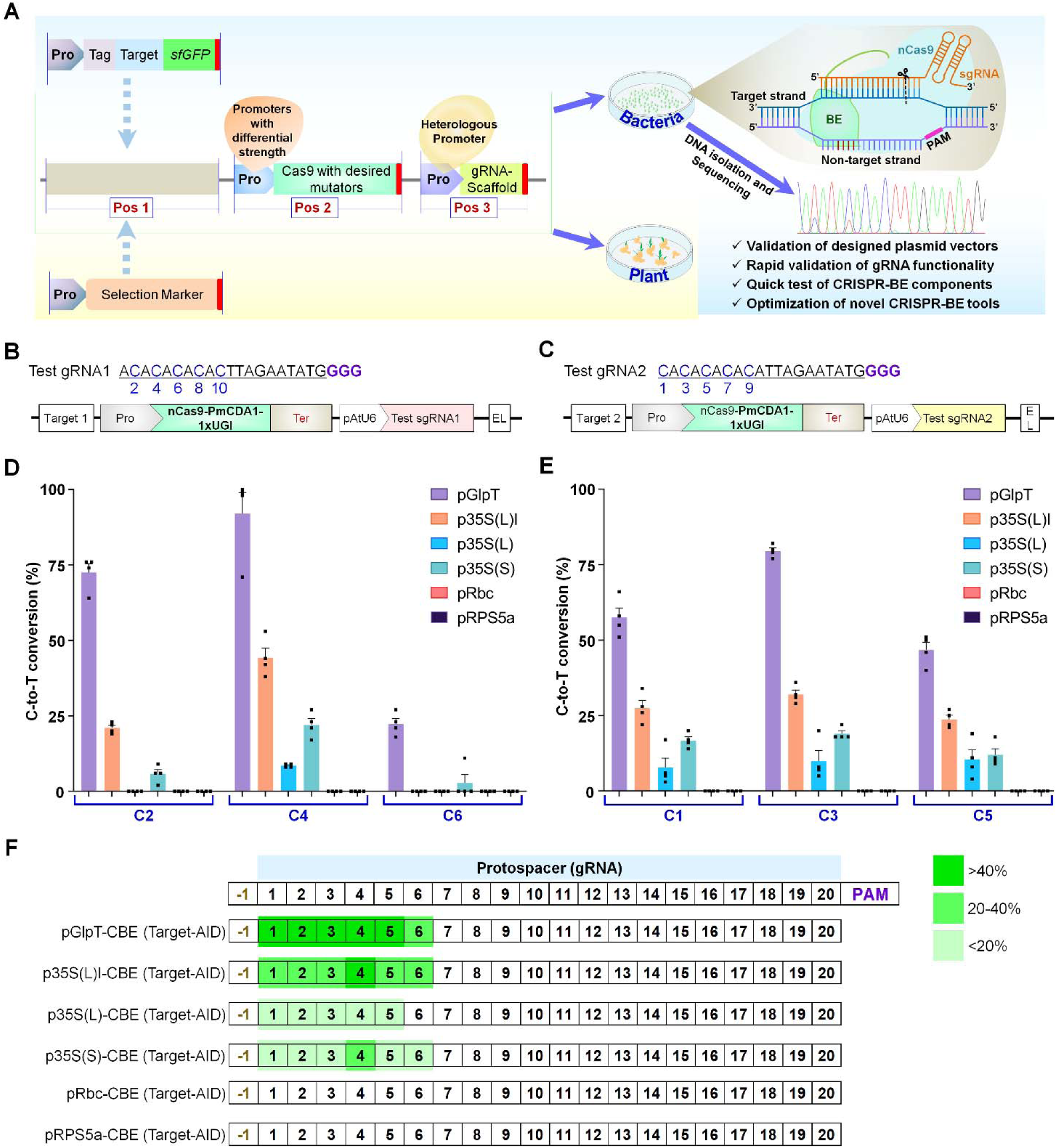
Evaluation of C-to-T conversion by PmCDA1-mediated cytosine base editor (CBE) driven by different promoters in *E. coli* cells. **(A)** Schematic representation of IRI-CCE (*in vivo* rapid investigation of CRISPR-BE components in *Escherichia coli*) platform. Promoters of differential-strength may enable the production of CRISPR-BE components (Cas9-fusions and sgRNAs) without toxic effects in *E. coli* allowing the testing of new BE tools, reliability of cloned plasmids, and editing activities of designed gRNAs. (**B-C)** Two independent gRNAs were tested containing alternate Cs at even (Test gRNA1, **B**) and odd (Test gRNA2, **C**) positions spanning from 1 to 10 in the 20 bp protospacer length. The numbers are assigned by counting the distal base as 1 from the protospacer adjacent motif (PAM), i.e., GGG. Schematic representation of the plasmid vectors showing a synthesized target region for gRNA and two transcriptional units (TUs) composing of nCas9 (D10A) fused with PmCDA1-1xUGI and AtU6 promoter-sgRNA unit. **(D-E)** PmCDA1-based C-to-T editing activities. Graph values show the mean percentage on the y-axis and the tested protospacer positions on the x-axis. The graph bar shows the mean of percentage values, and error bars indicate the standard error of the mean (mean ±s.e.m.) of four independent biological replicates. Dots indicate the individual biological replicates. (**F)** Summary of C-to-T editing activity windows of tested gRNAs for PmCDA1-CBE expressed by different promoters. Dark green zone (>40%) indicates the ideal editing window with higher editing efficiencies, and green (20-40%) or light green (<20%) zone regions indicate lesser C-to-T conversion.

After transforming individual constructs in 10-beta *E. coli*, single colonies were cultured for plasmid isolation and target-region sequencing. pEc1-driven nCas9-PmCDA1 shown poor transformation efficiency, probably attributable to lethal effect on bacterial growth. Base conversion (C-to-T) frequency of nCas9-PmCDA1 was found to be 100% in the case of pGlpT, p35S(L)I, p35S(L), and p35S(S) with variable efficiency in the editing window ranging from 1 to 6 position (**Figure 2D-E**). In comparison, pRbc and pRPS5a clones shown no C-to-T editing in the target region, supporting the absence of detectable sfGFP expression in **Figure 1**C. Calculated editing efficiency showed a variable percentage of C-to-T editing (**Figure 2D-F**), implying that after transforming the plasmid vectors into competent cells, the grown individual clones were indeed mixed populations. Similar outcomes for base editing were previously reported in other bacteria, like *Corynebacterium glutamicum* (Wang et al., 2018), *Clostridium beijerinckii* (Li et al., 2019), *Staphylococcus aureus* (Zhang et al., 2020), and *Agrobacterium* sp. (Rodrigues et al., 2021). For pGlpT, four colonies showed an average of 92% editing at targeted C4 in Test gRNA1 and 79.5% editing at targeted C5 in Test gRNA2 (**Figure 2D-E**). A variable range of editing efficiencies was observed for each C position in nCas9-PmCDA1 driven by different promoters, possibly due to differential expression of CRISPR-BE reagents. Consequently, the optimized method including differential-strength promoters for expression of CRISPR components was named IRI-CCE, an *In vivo* <u>R</u>apid Investigation of <u>C</u>RISPR <u>C</u>omponents in *<u>E</u>. coli*.

### 3.3 Assessing base editing activities on chromosomal targets in *E. coli* genome

Next, we targeted *E. coli* genes to evaluate the BE efficacy and potential toxicity mediated by differential-strength promoters in the genomic context of bacteria. Five different gRNAs were chosen for editing three genes, namely *galK, rpoB*, and *rppH* (**Figure 3**). The pGlpT and p35S(L)I-driven Target-AID (nCas9-PmCDA1) efficiently edited the available Cs in the editing window of all the tested sites without toxic effects on cell survival. Because the editing efficiency of BE versions may differ due to distinct genetic backgrounds in different *E. coli* strains, we also examined the base conversion efficiency in three more strains, including DH5α, DB3.1, and BL21(DE3). Irrespective of the genetic background of strain, we found comparable C-to-T mutations in tested gRNAs in all the three bacterial strains (data not shown), suggesting the broader applicability of the IRI-CCE platform across different *E. coli* strains.

**Figure 3.**
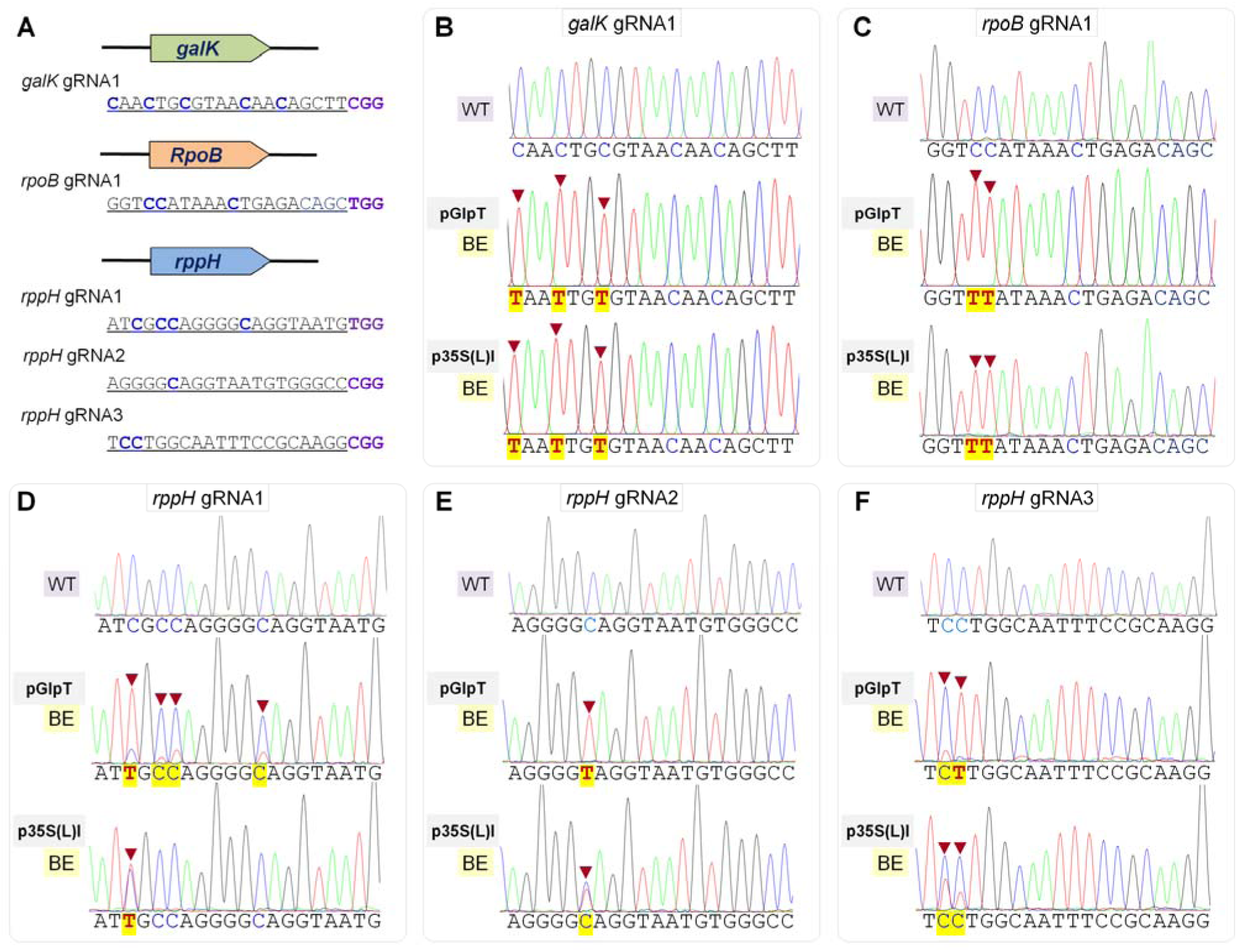
Differential-strength promoter-driven C-to-T editing activities by PmCDA1 in *E. coli* genome. **(A)** Designed gRNAs and PAM site (violet) for targeting three genes in the *E. coli* genome. Out of five gRNAs, 1, 1, and 3 were used for targeting *galK, rpoB*, and *rppH*, respectively. All the cytosine nucleotides located in the protospacer are highlighted in blue (bold). **(B-F)** Sequence alignment of the PmCDA1-edited mutants. The pGlpT and p35S(L)I-driven Target-AID efficiently edited the available Cs in the editing window of all the tested sites. Notably, in the p35S(L)I-driven Target-AID system, higher C-to-T activity was observed in *galK* and *rpoB*, which might be attributable to the limited number of available Cs in the editing window compared to multicopy plasmid-encoded targets.

### 3.4 Investigation of broad-range CBEs using IRI-CCE platform

Base editors with broader editing windows are generally applied for mutagenesis of a user-defined region. The editing windows vary according to BE type, target site, sequence context, and experimental conditions (Huang et al., 2021). Several latest BE tools are not yet characterized for *E. coli* use. To characterize the editing features in *E. coli*, two CBE types (evoCDA1 and A3A) were tested in the IRI-CCE platform with distinctive features (different range of editing window and no sequence context-preference). In original reports, editing windows for evoCDA1 and A3A positioned from 1 to 14 (Thuronyi et al., 2019) and 1 to 17 (Zong et al., 2018). A core-editing region was observed between 3 to 12 positions for evoCDA1 in human cells and 3 to 9 positions for A3A in plants. To examine C-to-T conversion in *E. coli*, in addition to two Test gRNAs, we designed Test gRNA3 by mimicking the sequence of TaVRN1-gRNA1 (**Figure 4A**) that comprises the Cs in the range of 1 to 17 bp (Zong et al., 2018).

**Figure 4.**
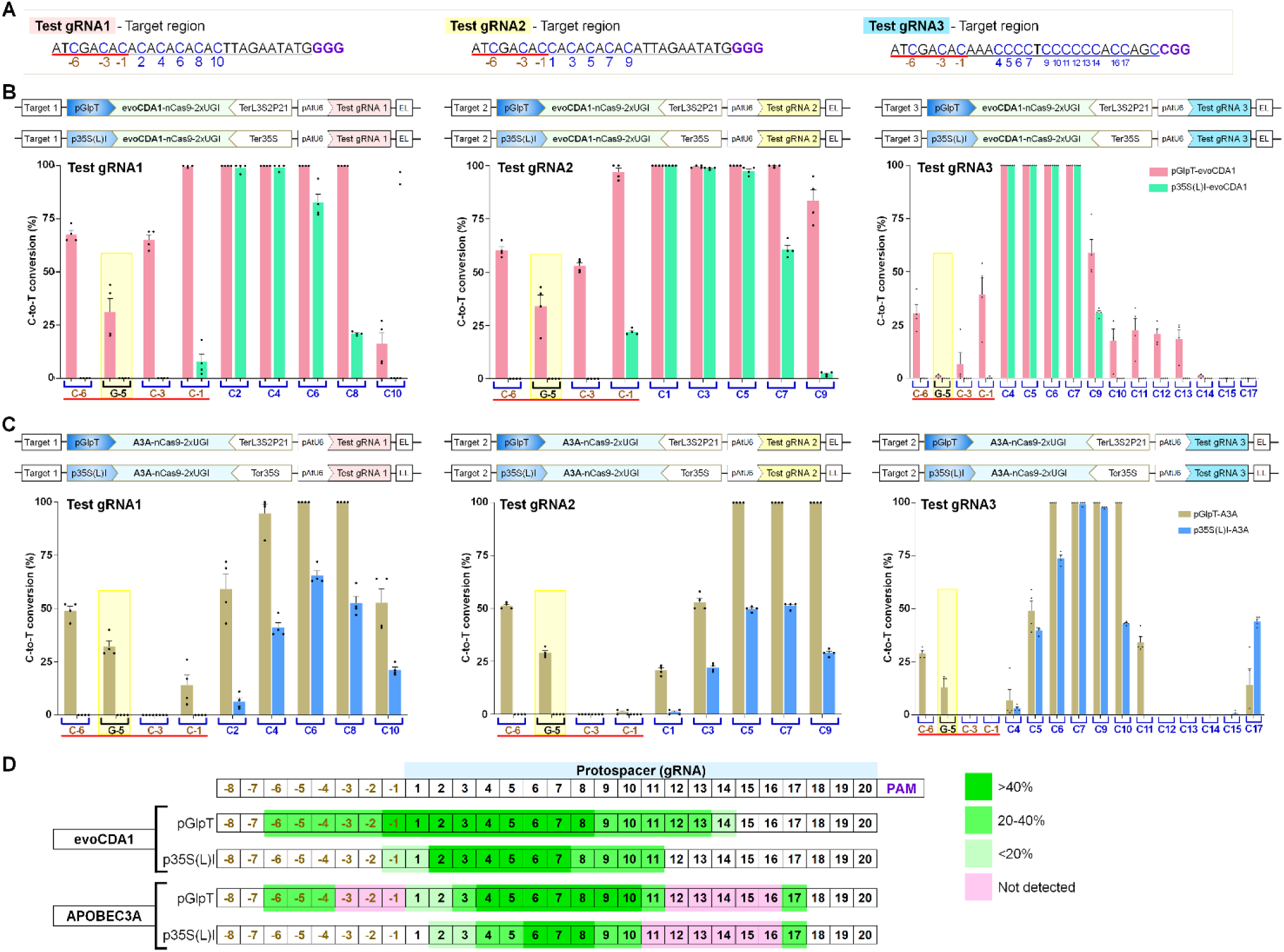
evoCDA1 and APOBEC3A expressed under the control of different promoters show expanded editing windows with efficient C-to-T editing. **(A)** Nucleotide sequences of three gRNAs (underlined in black) along with additional N-terminal regions (underlined in red). The numbers are assigned by counting the distal base as 1 from the protospacer adjacent motif (PAM), i.e., GGG. (**B-C)** evoCDA1 (**B**) and APOBEC3A (A3A) (**C**) driven C-to-T editing activities for three substrates targeted in three independent experiments. The designed BE constructs are represented on the upper side and obtained results depicted in bar graphs. G-5 position (yellow shade) data showed G•C to A•T conversion, suggesting editing happened in the target strand. Graph values show the mean percentage on the y-axis and the tested protospacer positions on the x-axis. The graph bar shows the mean of percentage values, and error bars indicate the standard error of the mean (mean ±s.e.m.) of four independent biological replicates. Dots indicate the individual biological replicates. (**D)** Summary of C-to-T editing activity windows of tested gRNAs for evoCDA1 and A3A expressed by different promoters. Dark green zone (>40%) indicates the ideal editing window with higher editing efficiencies, and green (20-40%) or light green (<20%) zone regions indicate lesser C-to-T conversion. The pink zone indicates the inaccessible regions for C-to-T editing.

Constitutive expression of CRISPR components using different promoters showed an expanded editing window for evoCDA1 and A3A, possibly owing to higher nCas9-BE expression (**Figure 4**). Among the tested CBEs, evoCDA1 showed the C-to-T conversion with the broadened editing window from -6 to 14 position with pGlpT and from -1 to 11 position with p35S(L)I (**Figure 4B**). The expression of nCas9-A3A using pGlpT and p35S(L)I promoter also displayed distinct editing windows ranging from -6 to 17 and 2 to 17, respectively (**Figure 4C**). For pGlpT-evoCDA1, C-to-T conversion at positions -1 to 9 was the highest (18%-100%), whereas, for pGlpT-A3A, C-to-T mutation at position 3 to 10 was the highest (43%-100%).

On the other hand, for p35S(L)I-evoCDA1, C-to-T conversion at positions 2 to 7 was the highest (57%-100%), whereas for p35S(L)I-A3A, C-to-T mutation at position 4 to 10 was the highest (38%-100%). Notably, G:C at position -5 was mutated to A:T in both the CBE types with moderate efficiency across all the three gRNAs (up to 43%), implying the C-to-T conversion in the target (non-deaminated) strand. The upstream (position -3 to -1) and middle (position 12 to 16) regions of target sites were not accessible to A3A deamination in the tested combinations of gRNAs and respective target sites (**Figure 4D**). Although direct comparison with earlier reports is not possible due to earlier mentioned parameters, these findings demonstrate the robust applicability of the IRI-CCE platform to investigate features of broad-range CBE tools.

### 3.5 ABE8e and ABE9e activities for A-to-G conversion in *E. coli*

Recently, evolved ABE variants, TadA-8e (ABE8e) and TadA-9e (ABE9e), have been characterized and shown improved A-to-G base conversion *in vitro*, human, and plants, respectively (Richter et al., 2020; Yan et al., 2021) but not yet systematically evaluated in bacteria. The gRNAs with the multiple As in a target region exhibited different A-to-G editing efficiencies for each position in *E. coli* and *S. aureus* (Zhang et al., 2020). Moreover, the editing window for ABE was also reported to vary with the target site, host species, genotype, and experimental conditions (Katti et al., 2020; Huang et al., 2021). We sought to investigate the ABE8e and ABE9e activity using two synthetic gRNAs (**Figure 5**). The editing window for ABE8e and ABE9e variants was reported as positions 4 to 8 in human cells (Richter et al., 2020) and 1 to 11 in plants (Yan et al., 2021), respectively. Editing levels of ABE8e and ABE9e showed significant increases in the case of pGlpT compared to that of p35S(L)I (**Figure 5B-D**).

**Figure 5.**
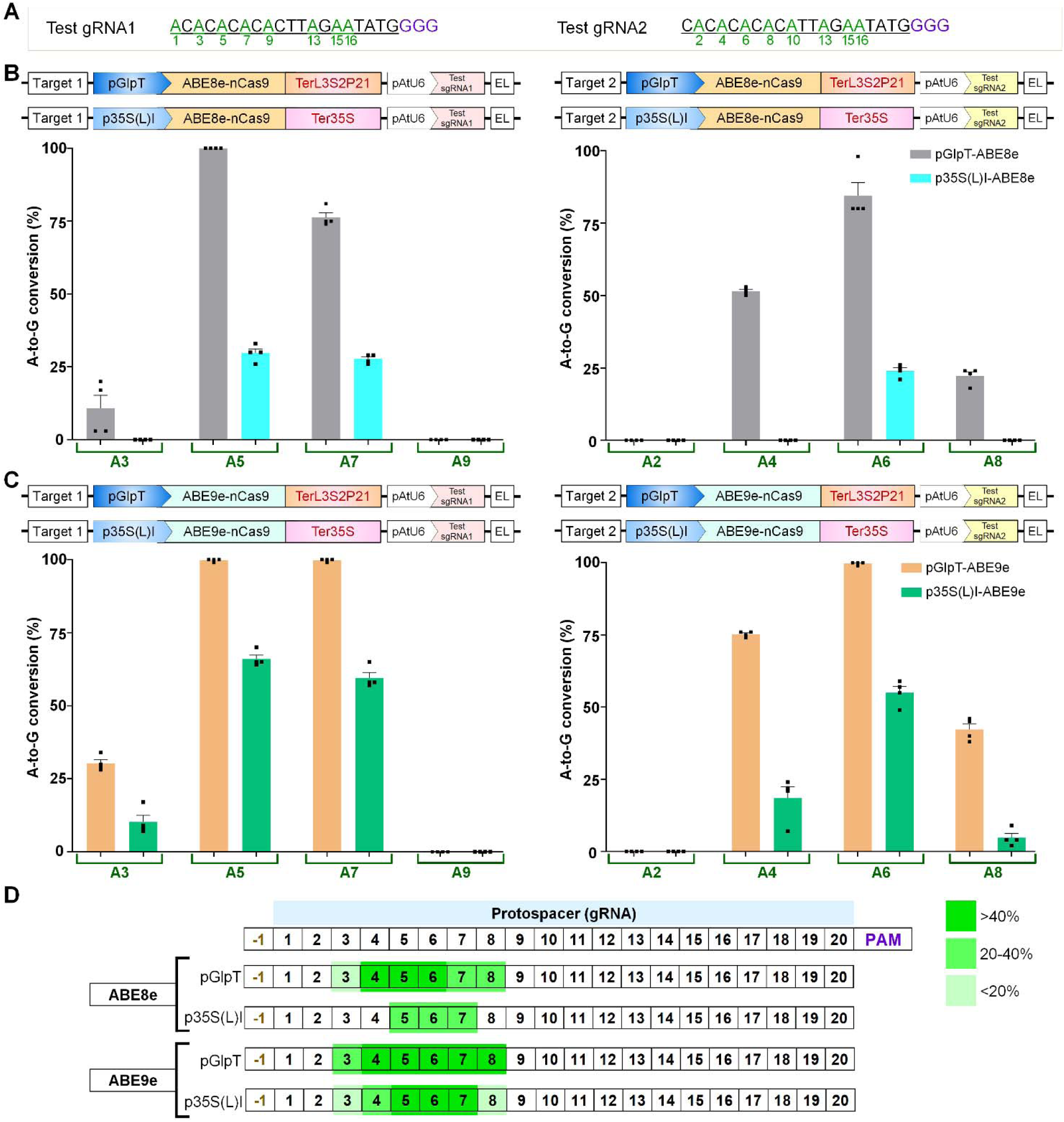
Adenine base editor variants (ABE8e and ABE9e) efficiently convert A-to-G in *E. coli* cells. **(A)** Nucleotide sequences of Test gRNA1 and 2 (underlined in black) showing available adenine (A) residues (green). **(B-C)** The A-to-G conversion efficiencies of ABE8e **(B)** and ABE9e **(C)** driven by pGlpT and p35S(L)I promoters in the target region for Test gRNA1 and Test gRNA2. The graph bar shows the mean of percentage values, and error bars indicate the standard error of the mean (mean ±s.e.m.) of four independent biological replicates. Dots indicate the individual biological replicates. (**D)** Summary of A-to-G editing windows of tested gRNAs for ABE8e and ABE9e.

**Figure 6.**
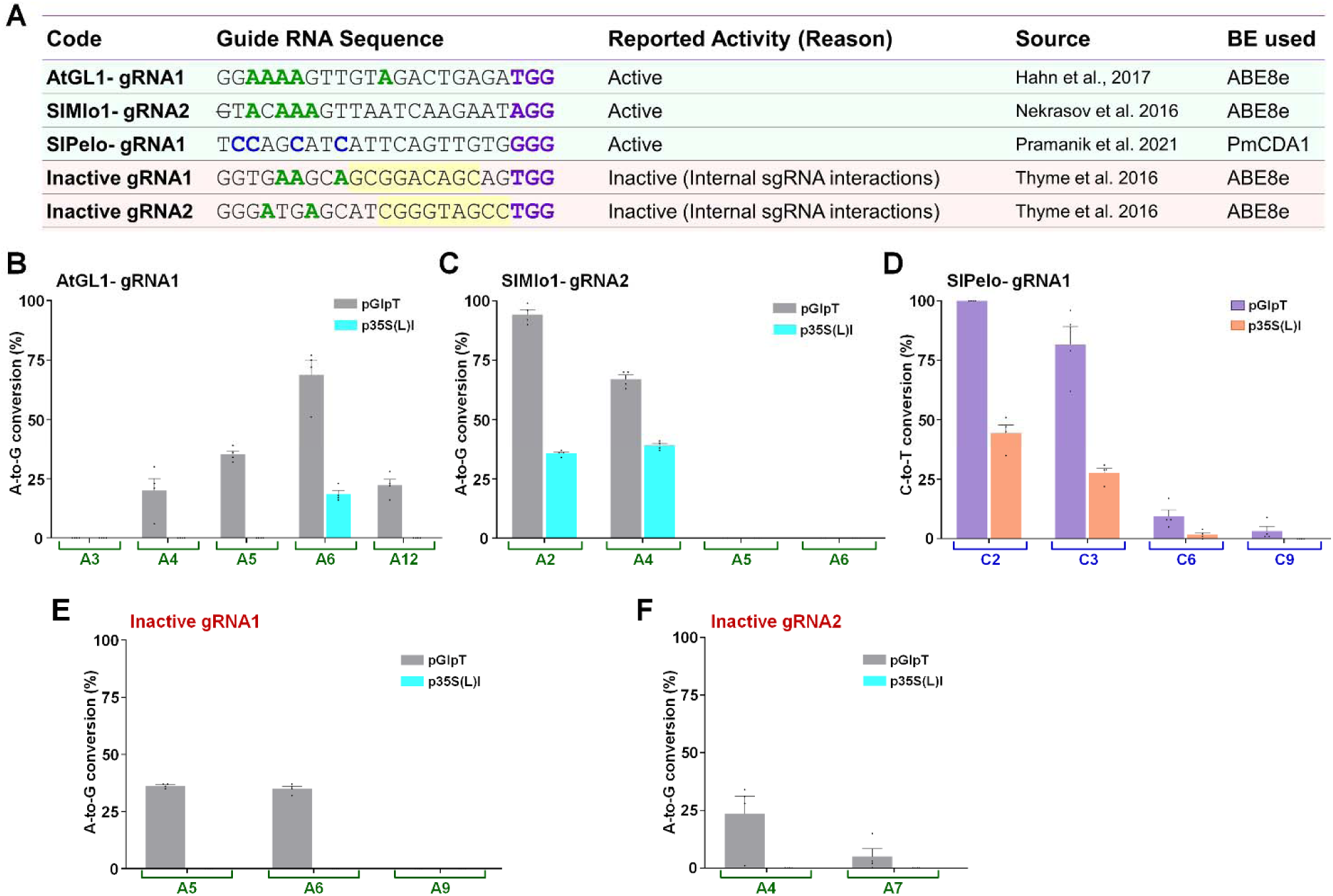
Screening of designed gRNAs in the IRI-CCE platform provides first-hand knowledge about the sgRNA functionality. **(A)** Set of gRNAs used for validating the potential use of IRI-CCE in gRNA screening. The base editor (BE) type was chosen depending on the availability of A or C in the editing window. Frequencies of base conversions by utilized BE types in target regions of AtGL1-gRNA1 (**B**), SlMlo1-gRNA2 (**C**), and SlPelo-gRNA1 (**D**), Inactive gRNA1 (**E**), and Inactive gRNA2 (**F**). The graph bar shows the mean of percentage values, and error bars indicate the standard error of the mean (mean ±s.e.m.) of four independent biological replicates. Dots indicate the individual biological replicates.

We observed the editing window of ABE8e from positions 3 to 8 and 5 to 7 for pGlpT and p35S(L)I, respectively (**Figure 5D**). While editing window of ABE9e for pGlpT and p35S(L)I ranged from positions 3 to 8 and 3 to 7, respectively. A-to-G conversion at positions 5 to 7 was the highest in the case of both the variants (ABE8e: 74%-100%; ABE9: 99-100%). For p35S(L)I-ABE8e, A-to-G mutation at positions 5 to 7 was the highest but in the lower range (21%-33%), and p35S(L)I-ABE9e shown a moderate editing activity (55-66%). These data suggest that the editing activity of ABE9e was more enriched than that of ABE8e in *E. coli*, and editing activity increases in a promoter strength-dependant manner.

### 3.6 IRI-CCE platform allows screening of functional gRNAs

Pre-screening of gRNAs before commencing GE studies is crucial in higher eukaryotes to avoid loss of time for gRNA screening, lower editing activities, and waste of resources. Given that the efficiencies predicted by *in silico* tools and the actual GE activities may differ significantly, *in vitro* or preferentially *in vivo* validation of chosen gRNAs is vital before the actual tests (Kim et al., 2021). In that scenario, the IRI-CCE platform may facilitate the simultaneous evaluation of designed bioparts and functionality of the selected gRNAs that enable the potential avoidance of incorrect plasmids and inefficient gRNAs. In some instances, gRNAs predicted as inactive by bioinformatic tools or design criteria were reported to work efficiently under *in vivo* conditions (Naim et al., 2020). On the other hand, even if *in silico* parameters are not promising for a particular set of gRNAs, one needs to choose those gRNAs for editing a specific region of targeted locus due to PAM constraints.

We analyzed sets of active and inactive gRNAs reported in the previous studies (**Figure 5A**). Based on the A or C availability in the canonical editing window, we opted for a particular BE type to edit the target sites of individual gRNAs. The target regions, including the PAM, were cloned using two BsmBI type IIS sites in the universal target-acceptor modules (**Supplementary Figures 5 and 6**). Firstly, the set of three active gRNAs (AtGL1-gRNA1 (Hahn et al., 2017), SlMlo1-gRNA2, and SlPelo-gRNA1 (Pramanik et al., 2021b) targeting three plant genomic targets showed higher nucleotide conversion efficiencies (**Figure 5B-D**) with differential-strength promoters as observed under *in planta* conditions.

Secondly, we analyzed ABE8e activities for two inactive gRNAs (Inactive gRNA1 and Inactive gRNA2) that displayed poorer performance (Thyme et al., 2016). The lower editing activity was attributed to internal sgRNA interactions that allow binding with Cas9, but it prevents target DNA recognition by the Cas9-sgRNA complex (**Supplementary Figure 7**). Besides, inactive gRNA competes with active gRNAs for binding to the Cas9 protein, thereby reducing the editing activities. We investigated the base conversion activity at DNA templates for both the gRNAs and discovered that both the sgRNAs led to no BE frequencies in the target DNA region with low-strength p35S(L)I (**Figure 5E-F**), consistent with the *in vivo* data for CRISPR/Cas editing (Thyme et al., 2016). Moreover, even a high-strength promoter (pGlpT)-driven ABE8e expression led to relatively lower BE activity (36% and 23.5% for Inactive gRNA1 and 2, respectively). Overall, these data indicate that the IRI-CCE platform is helpful to distinguish the functional gRNAs, although with limited information (elaborated in the discussion section). Taken together, the IRI-CCE platform could provide first-hand knowledge of *in vivo* gRNA activity and help to reduce the number of gRNAs one needs to use in higher eukaryotic systems.

## 4 Discussion

The several features of *E. coli* make it a model microorganism in molecular biology. It includes a short life cycle, simple genetic manipulations, low cost, and in-depth knowledge of the genome (Jia and Jeon, 2016). In the current study, we developed an *E. coli*-based IRI-CCE platform to investigate two key factors: cloned BE bioparts for modular cloning and gRNA functionality with several advantages. First, IRI-CCE enables the rapid validation of designed BE bioparts for further experiments. Secondly, modular bioparts are exchangeable for cloning and investigating different CRISPR-based BE tools in bacteria and plants, including deaminases, nCas9, heterologous promoters (p35S and pAtU6), accessory components like UGI, etc. Third, quick verification of gRNA functionality may help to reduce the burden of gRNAs screening in eukaryotes (**Figure 2A**). Also, the approach reported here could be readily scaled to many constructs and different BE types and evolved versions.

The attempt to analyze the BE activity of bacterial (pJ23119) promoter-driven gRNA expression was unsuccessful due to the toxicity caused by BE components. Likewise, the pEc1-mediated CBE test also showed poor transformation efficiency with a mix of incorrect clones. The toxicity of BE components is earlier reported in bacteria (Banno et al., 2018; Rodrigues et al., 2021). When nCas9-PmCDA1 was expressed together with pJ23119-sgRNA, no correct clones were obtained in *E. coli*, possibly due to toxic effects of constitutively-expressed BE components (Banno et al., 2018). The cloning of dCas9-PmCDA1 with pJ23119-sgRNA showed the wrong clones in *E. coli* (Banno et al., 2018) and *Agrobacterium* sp. (Rodrigues et al., 2021), suggesting cells cannot retain the functional dCas9-PmCDA1-expressing plasmids. A high rate of SSBs induced by nCas9 during pJ23119-mediated sgRNA production seems difficult to be restored by the bacterial system, ultimately losing the plasmid. Although dCas9 does not introduce SSBs, earlier dCas9-PmCDA1 data suggested that the surplus amount of UGI compromises the genome integrity in CBE types (Banno et al., 2018).

Previous studies reported that the use of protein degradation (LVA) tag fusion with dCas9-Target-AID reduced the toxicity to *E. coli* cells (Banno et al., 2018). By producing the optimal amount of BE and sgRNA might reduce the toxicity to a tolerable level for a bacterial cell as observed in nCas9-BE together with pAtU6-driven sgRNA expression. As described in our study, the combination of pGlpT-nCas9-BE or p35S(L)-nCas9-BE with pAtU6-sgRNA expression units proved to be the most appropriate to edit the target site in the same plasmid vector or *E. coli* genome. Therefore, fine-tuning the expression of BEs and sgRNAs by using the optimal combination of promoters is a crucial factor in minimizing the toxic effects in the IRI-CCE platform.

Deaminase interaction with Cas protein and target DNA region alters the location and size of the editing window (Cheng et al., 2019). The indispensable role of the endogenous DNA repair mechanism is a decisive factor that may produce variable BE results in different organisms (Kurt et al., 2021). Nevertheless, we observed significant differences between the targeting competencies of three CBEs owing to the differential strength of promoters used for Cas-BE expression (**Supplementary Figure 8**). The evoCDA1 is an evolved variant of PmCDA1 (Thuronyi et al., 2019) that showed higher catalytic activity and expanded editing window than its wild-type counterpart. We observed higher C-to-T editing in the expanded window up to position -6 that was not detected in the previous work for evoCDA1 or A3A.

The comparison between the editing windows revealed that the accessibility to the target site was different for evoCDA1 and A3A, possibly due to differences in R-loop formations and editing competencies. Some Cs in the editing window were not accessible to A3A deamination. These observations highlight the importance of considering BE type and promoter strength that best suit the expected outcome. Also, temperature affects the enzymatic activities of CRISPR components, thereby influencing the GE outcome (Pramanik et al., 2021a). Mainly, the performance of Cas enzymes is best at 37°C. So, besides the promoter activity and evolved variants, better editing efficacy in IRI-CCE may resulted from the favorable temperatures for enzymatic activities.

Some of the gRNA-related factors like nCas9-BE-sgRNA complex formation or stability and target DNA-binding would be identified in the IRI-CCE platform. However, more efforts are needed to understand uncovered aspects. For example, the editing efficiency of each nCas9-BE-sgRNA complex differs significantly, depending on the target locus (Pramanik et al., 2021b). In higher eukaryotes, editing differences arise from the sequence bias in DNA repair machinery and target accessibility (Isaac et al., 2016). Also, the bigger genome size of eukaryotes influences the dynamics of target search. Therefore, IRI-CCE is missing those features common to eukaryotic cells and cannot be verified in the current platform. While our data show the promising way to confirm the biopart modules for CRISPR-based BE work, there are multiple opportunities to expand the IRI-CCE utility further. For example, the design and adoption of new dual BEs by combining different variants of ABE and CBE could be interesting for simultaneous conversion of A-to-G and C-to-T, respectively.

## 5 Conclusions

In summary, nCas9-BE components expressed by promoters of different strengths led to the establishment of the non-toxic IRI-CCE platform for two major applications: investigation of cloned CRISPR-BE plasmids and to know gRNA functionality. IRI-CCE analyses showed the variable length of the editing window for specific BE types. Each independent BE type has its specific editing pattern for a given target site and promoter type. Because of its easy cloning and heterologous applicability, the IRI-CCE platform may be suited for rapid evaluation of cloned bioparts in modular cloning of BE studies and applicable to a broader range of *E. coli* strains being used for cloning. This platform facilitates the verification of gRNA-related factors such as cloning, sgRNA production, Cas9-sgRNA complex formation, and DNA recognition. Through the applications described here and through further improvements, IRI-CCE can be widely applicable for the characterization of BE components and rapid assessment of newer CRISPR-based BE technologies.

## 7 Author Contributions

J.Y.-K. and R.M.S. conceived this study and designed the research. R.M.S. designed constructs and performed experiments. R.M.S. and D.P. performed analyses. R.M.S. and D.P. wrote the paper and analyzed the data. J.Y.-K. supervised the research activities. All authors contributed to discussions and manuscript review.

## 10 Supplementary Material

**Supplementary Figure 1**. Promoter sequences from different organisms used in **Figure** 1.

**Supplementary Figure 2**. Evaluation of promoter activities at the transcriptional level in *E. coli*.

**Supplementary Figure 3**. Promoter sequences used for sgRNA expression in different organisms.

**Supplementary Figure 4**. Promoter sequence alignment used for sgRNA expression in bacteria (J23119) and plant (Arabidopsis U6).

**Supplementary Figure 5**. The scheme followed for cloning the target regions as DNA templates for gRNA binding and subsequently CRISPR editing.

**Supplementary Figure 6**. Approaches analyzed for the establishment of IRI-CCE platform.

**Supplementary Figure 7**. Features of inactive gRNAs from **Figure** 5.

**Supplementary Figure 8**. Editing windows for CBEs (PmCDA1, evoCDA1, APOBEC3A) and ABE reported in the present study expressed under the promoters of different strengths.

**Supplementary Table 1**. DNA sequences of CRISPR and BE components.

**Supplementary Table 2**. *Escherichia coli* strains used in present study.

**Supplementary Table 3**. Primer sequences used for cloning and sequencing are summarized.

**Supplementary Table 4**. Plasmids used in present work.

## 11 Data Availability Statement

The raw data supporting the conclusions of this article will be made available by the authors, without undue reservation.

