## Supplementary File 1 for "IRI-CCE: A Platform for Evaluating CRISPR-based Base Editing Tools and Its Components"

| | **Content** | **Page** |
|---|---|---|
| 1 | Supplementary Fig. 1 Promoter sequences from different organisms used in Fig. 1. | 3 |
| 2 | Supplementary Fig. 2 Evaluation of promoter activities at the transcriptional level in *E. coli*. | 4 |
| 3 | Supplementary Fig. 3 Promoter sequences used for sgRNA expression in different organisms. | 5 |
| 4 | Supplementary Fig. 4 Promoter sequence alignment used for sgRNA expression in bacteria (J23119) and plant (Arabidopsis U6). | 6 |
| 5 | Supplementary Fig. 5. The scheme followed for cloning the target regions as DNA templates for gRNA binding and subsequently CRISPR-BE editing. | 7 |
| 6 | Supplementary Fig. 6. Approaches analyzed for the establishment of IRI-CCE platform. | 8 |
| 7 | Supplementary Fig. 8. Features of inactive gRNAs from Fig. 5. | 9 |
| 8 | Supplementary Fig. 9. Editing windows for CBEs (PmCDA1, evoCDA1, APOBEC3A) and ABE reported in the present study expressed under the promoters of different strengths. | 10 |
| 9 | Supplementary Table 1. DNA sequences of CRISPR and BE components. | 11 |
| 10 | Supplementary Table 2. *Escherichia coli* strains used in the present study. | 12 |
| 11 | Supplementary Table 3. Primer sequences used for cloning and sequencing are summarized. | 16 |
| 12 | Supplementary Table 4. Plasmids used in the present work. | 18 |
| 13 | **References** | 21 |

**pEc1** (SJM901+RBSTL2)

TTTACAGCTAGCTCAGTCCTAGGTATAATGCTAGCAGATCTAATAATTTTGTTTAACTTTGGGAGGATA

**pGlpT** (BBa_J72163 GlpT+RBS)

GAAAGTGAAACGTGATTTCATGCGTCATTTTGAACATTTTGTAAATCTTATTTAATAATGTGTGCGGCAATTCACATTTAATTTATGAATGTTTTCTTAACATCGCGGCAACTCAA
GAAACGGCAGGTTCGGATCTTAGCTACTAGAGAAAGAGGAGAAATACTAG

**p35S(L)I** [CaMV 35S(Long)+AtUBQ10 Intron I +TMV5'U]

GAATTCCAATCCCACAAAAATCTGAGCTTAACAGCACAGTTGCTCCTCTCAGAGCAGAATCGGGTATTCAACACCCTCATATCAACTACTACGTTGTGTATAACGGTCCACATGCC
GGTATATACGATGACTGGGGTTGTACAAAGGCGGCAACAAACGGCGTTCCCGGAGTTGCACACAAGAAATTTGCCACTATTACAGAGGCAAGAGCAGCAGCTGACGCGTACACAAC
AAGTCAGCAAACAGACAGGTTGAACTTCATCCCCAAAGGAGAAGCTCAACTCAAGCCCAAGAGCTTTGCTAAGGCCCTAACAAGCCCACCAAAGCAAAAAGCCCACTGGCTCACGC
TAGGAACCAAAAGGCCCAGCAGTGATCCAGCCCCAAAAGAGATCTCCTTTGCCCCGGAGATTACAATGGACGATTTCCTCTATCTTTACGATCTAGGAAGGAAGTTCGAAGGTGAA
GGTGACGACACTATGTTCACCACTGATAATGAGAAGGTTAGCCTCTTCAATTTCAGAAAGAATGCTGACCCACAGATGGTTAGAGAGGCCTACGCAGCAAGTCTCATCAAGACGAT
CTACCCGAGTAACAATCTCCAGGAGATCAAATACCTTCCCAAGAAGGTTAAAGATGCAGTCAAAAGATTCAGGACTAATTGCATCAAGAACACAGAGAAAGACATATTTCTCAAGA
TCAGAAGTACTATTCCAGTATGGACGATTCAAGGCTTGCTTCATAAACCAAGGCAAGTAATAGAGATTGGAGTCTCTAAAAAGGTAGTTCCTACTGAATCTAAGGCCATGCATGGA
GTCTAAGATTCAAATCGAGGATCTAACAGAACTCGCCGTCAAGACTGGCGAACAGTTCATACAGAGTCTTTTACGACTCAATGACAAGAAGAAAATCTTCGTCAACATGGTGGAGC
ACGACACTCTGGTCTACTCCAAAAATGTCAAAGATACAGTCTCAGAAGATCAAAGGGCTATTGAGACTTTTCAACAAAGGATAATTTCGGGAAACCTCCTCGGATTCCATTGCCCA
GCTATCTGTCACTTCATCGAAAGGACAGTAGAAAAGGAAGGTGGCTCCTACAAATGCCATCATTGCGATAAAGGAAAGGCTATCATTCAAGATCTCTCTGCCGACAGTGGTCCCAA
AGATGGACCCCCACCCACGAGGAGCATCGTGGAAAAAGAAGAGGTTCCAACCACGTCTACAAAGCAAGTGGATTGATGTGACATCTCCACTGACGTAAGGGATGACGCACAATCCC
ACTATCCTTCGCAAGACCCTTCCTCTATATAAGGAAGTTCATTTCATTTGGAGAGGACACGCTCGAGTATAAGGTAAATTTCTGTGTTCCTTATTCTCTCAAAATCTTCGATTTTG
TTTTCGTTCGATCCCAATTTCGTATATGTTCTTTGGTTTAGATTCTGTTAATCTTAGATCGAAGATGATTTTCTGGGTTTGATCGTTAGATATCATCTTAATTCTCGATTAGGGTT
TCATAGATATCATCCGATTTGTTCAAATAATTTGAGTTTTGTCGAATAATTACTCTTCGATTTGTGATTTCTATCTAGATCTGGTGTTAGTTTCTAGTTTGTGCGATCGAATTTGT
CGATTAATCTGAGTTTTTCTGATTAACAGGAGCTCATTTTTACAACAATTACCAACAACAACAAACAACAAACAACATTACAATTACATTTACAATTATCGATAC

**p35S(L)** [CaMV 35S(Long)+TMV5'U]

GAATTCCAATCCCACAAAAATCTGAGCTTAACAGCACAGTTGCTCCTCTCAGAGCAGAATCGGGTATTCAACACCCTCATATCAACTACTACGTTGTGTATAACGGTCCACATGCC
GGTATATACGATGACTGGGGTTGTACAAAGGCGGCAACAAACGGCGTTCCCGGAGTTGCACACAAGAAATTTGCCACTATTACAGAGGCAAGAGCAGCAGCTGACGCGTACACAAC
AAGTCAGCAAACAGACAGGTTGAACTTCATCCCCAAAGGAGAAGCTCAACTCAAGCCCAAGAGCTTTGCTAAGGCCCTAACAAGCCCACCAAAGCAAAAAGCCCACTGGCTCACGC
TAGGAACCAAAAGGCCCAGCAGTGATCCAGCCCCAAAAGAGATCTCCTTTGCCCCGGAGATTACAATGGACGATTTCCTCTATCTTTACGATCTAGGAAGGAAGTTCGAAGGTGAA
GGTGACGACACTATGTTCACCACTGATAATGAGAAGGTTAGCCTCTTCAATTTCAGAAAGAATGCTGACCCACAGATGGTTAGAGAGGCCTACGCAGCAAGTCTCATCAAGACGAT
CTACCCGAGTAACAATCTCCAGGAGATCAAATACCTTCCCAAGAAGGTTAAAGATGCAGTCAAAAGATTCAGGACTAATTGCATCAAGAACACAGAGAAAGACATATTTCTCAAGA
TCAGAAGTACTATTCCAGTATGGACGATTCAAGGCTTGCTTCATAAACCAAGGCAAGTAATAGAGATTGGAGTCTCTAAAAAGGTAGTTCCTACTGAATCTAAGGCCATGCATGGA
GTCTAAGATTCAAATCGAGGATCTAACAGAACTCGCCGTCAAGACTGGCGAACAGTTCATACAGAGTCTTTTACGACTCAATGACAAGAAGAAAATCTTCGTCAACATGGTGGAGC
ACGACACTCTGGTCTACTCCAAAAATGTCAAAGATACAGTCTCAGAAGATCAAAGGGCTATTGAGACTTTTCAACAAAGGATAATTTCGGGAAACCTCCTCGGATTCCATTGCCCA
GCTATCTGTCACTTCATCGAAAGGACAGTAGAAAAGGAAGGTGGCTCCTACAAATGCCATCATTGCGATAAAGGAAAGGCTATCATTCAAGATCTCTCTGCCGACAGTGGTCCCAA
AGATGGACCCCCACCCACGAGGAGCATCGTGGAAAAAGAAGAGGTTCCAACCACGTCTACAAAGCAAGTGGATTGATGTGACATCTCCACTGACGTAAGGGATGACGCACAATCCC
ACTATCCTTCGCAAGACCCTTCCTCTATATAAGGAAGTTCATTTCATTTGGAGAGGACACGCTCGAGTATAAGAGCTCATTTTTACAACAATTACCAACAACAACAAACAACAAAC
AACATTACAATTACATTTACAATTATCGATAC

**p35S(S)** [CaMV 35S(Short)+TMV5'U]

GTCAACATGGTGGAGCACGACACTCTGGTCTACTCCAAAAATGTCAAAGATACAGTCTCAGAAGATCAAAGGGCTATTGAGACTTTTCAACAAAGGATAATTTCGGGAAACCTCCT
CGGATTCCATTGCCCAGCTATCTGTCACTTCATCGAAAGGACAGTAGAAAAGGAAGGTGGCTCCTACAAATGCCATCATTGCGATAAAGGAAAGGCTATCATTCAAGATCTCTCTG
CCGACAGTGGTCCCAAAGATGGACCCCCACCCACGAGGAGCATCGTGGAAAAAGAAGAGGTTCCAACCACGTCTACAAAGCAAGTGGATTGATGTGACATCTCCACTGACGTAAGG
GATGACGCACAATCCCACTATCCTTCGCAAGACCCTTCCTCTATATAAGGAAGTTCATTTCATTTGGAGAGGACACGCTCGAGTATAAGAGCTCATTTTTACAACAATTACCAACA
ACAACAAACAACAAACAACATTACAATTACATTTACAATTATCGATAC

**pRbc** (AtRbcS2B)

GCTTAAGTGGGCCGTAAGTGCAATTGCGTTGTGCTCCAAAAGGAACGGTGCCGTGGGTTCCTCTTGTTCATCAGAAATATATTAATTAGCCGTAAAACCTGAAAATTCACAAGCAT
TTGGATTGTTTTCTAATTTAATATCCATTATGTGACTAAAAGTTCTAGTGATCGTACATACTACATAGAAAATAATAACACAAAATACTAGTTTACATTTCCCAATTAAAAACCAT
TTTGAATGAACTCTGTCTGATTTAATTATACTTTTAAAATGTGGGATGAATTCAAAGATTATACTTATATTCTTATTATTTAAGATTATCAAGTGGAAAAATAAAAATATGAATGT
GTTAATATAAGGTAATAGAAATTTAATCATTTTTTTAATCTATATGTAAAAAGTATTTAACCGATATCTACAATTTGACGCCTCCCAATTGAAAGGAGCCAAAAGCAACCGATCAA
GTGCAGACCAGTAGCCATACACATTCACTCCTACCCTTACATGAGAAAGATAAGATTATGGAGTTTTCTGCCACGTGATCTTATCCTAGTGGTCCAAATCGATAAGGGTGTCAACA
CCTTTCCTTAATCCTGTGGCAATTAACGACGTTATCATGAATTATGGCCCCTTTGATCATTAGGGCTAGTTGCCTCTAGCGGTTCCCACTATATAAAGATGACAAAACCAACAGAC
AAACAAGTAAGTAAGAGAAAAACCAAAAGAAGAAGAGAAACAACAAGAAGAAGT

**pRPS5a**

GATCCCTCAACTTTTGATTCGCTATTTGCAGTGCACCTGTGGCGTTCATCACATCTTTTGTGACACTGTTTGCACTGGTCATTGCTATTACAAAGGACCTTCCTGATGTTGAAGGA
GATCGAAAGTAAGTAACTGCACGCATAACCATTTTCTTTCCGCTCTTTGGCTCAATCCATTTGACAGTCAAAGACAATGTTTAACCAGCTCCGTTTGATATATTGTCTTTATGTGT
TTGTTCAAGCATGTTTAGTTAATCATGCCTTTGATTGATCTTGAATAGGTTCCAAATATCAACCCTGGCAACAAAACTTGGAGTGAGAAACATTGCATTCCTCGGTTCTGGACTTC
TGCTAGTAAATTATGTTTCAGCCATATCACTAGCTTTCTACATGCCTCAGGTGAATTCATCTATTTCCGTCTTAACTATTTCGGTTAATTAAAGCACGAACACCATTACTGCATGT
AGAAGCTTGATAAACTATCGCCACCAATTTATTTTTGTTGCGATATTGTTACTTTCCTCAGTATGCAGCTTTGAAAAGACCAACCCTCTTATCCTTTAACAATGAACAGGTTTTTA
GAGGTAGCTTGATGATTCCTGCACATGTGATCTTGGCTTCAGGCTTAATTTTCCAGGTAAAGCATTATGAGATACTCTTATATCTCTTACATACTTTTGAGATAATGCACAAGAAC
TTCATAACTATATGCTTTAGTTTCTGCATTTGACACTGCCAAATTCATTAATCTCTAATATCTTTGTTGTTGATCTTTGGTAGACATGGGTACTAGAAAAAGCAAACTACACCAAG
GTAAAATACTTTTGTACAAACATAAACTCGTTATCACGGAACATCAATGGAGTGTATATCTAACGGAGTGTAGAAACATTTGATTATTGCAGGAAGCTATCTCAGGATATTATCGG
TTTATATGGAATCTCTTCTACGCAGAGTATCTGTTATTCCCCTTCCTCTAGCTTTCAATTTCATGGTGAGGATATGCAGTTTTCTTTGTATATCATTCTTCTTCTTCTTTGTAGCT
TGGAGTCAAAATCGGTTCCTTCATGTACATACATCAAGGATATGTCCTTCTGAATTTTTATATCTTGCAATAAAAATGCTTGTACCAATTGAAACACCAGCTTTTTGAGTTCTATG
ATCACTGACTTGGTTCTAACCAAAAAAAAAAAAATGTTTAATTTACATATCTAAAAGTAGGTTTAGGGAAACCTAAACAGTAAAATATTTGTATATTATTCGAATTTCACTCATCAT
AAAAACTTAAATTGCACCATAAAATTTTGTTTTACTATTAATGATGTAATTTGTGTAACTTAAGATAAAAATAATATTCCGTAAGTTAACCGGCTAAAACCACGTATAAACCAGGG
AACCTGTTAAACCGGTTCTTTACTGGATAAAGAAATGAAAGCCCATGTAGACAGCTCCATTAGAGCCCAAACCCTAAATTTCTCATCTATATAAAAGGAGTGACATTAGGGTTTTT
GTTCGTCCTCTTAAAGCTTCTCGTTTTCTCTGCCGTCTCTCTCATTCGCGCGACGCAAACGATCTTCAGGTGATCTTCTTTCTCCAAATCCTCTCTCATAACTCTGATTTCGTACT
TGTGTATTTGAGCTCACGCTCTGTTTCTCTCACCACAGCC

**Supplementary Figure 1. Promoter sequences from different organisms used in Figure 1.**

Additional features like ribosome binding sites (RBSs) in bacterial promoters are underlined. The intron region from *AtUbi10* in p35S(L)I is highlighted in violet, and the TMV 5U leader sequence is depicted in blue.

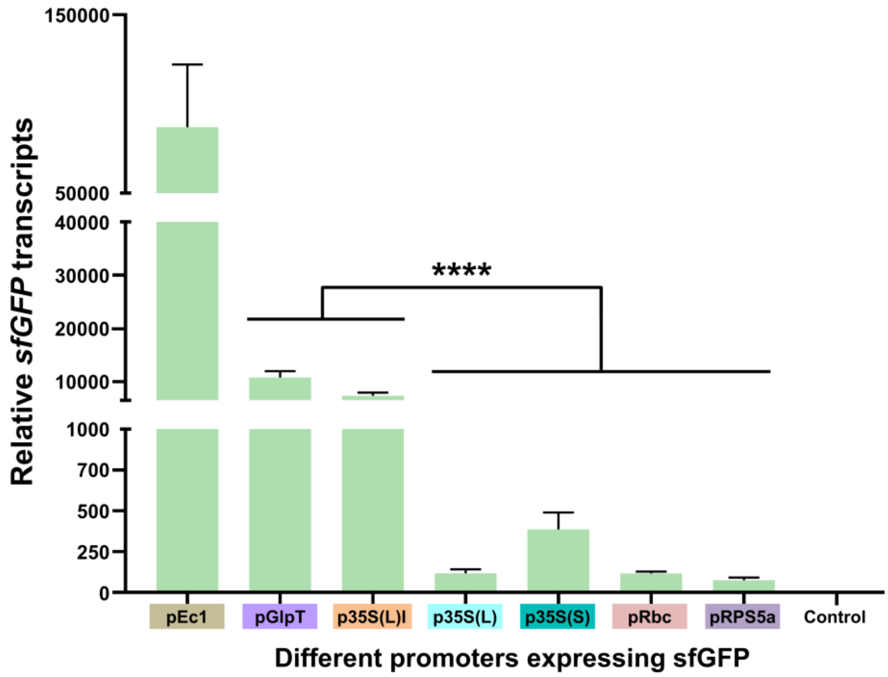


**Supplementary Figure 2. Evaluation of promoter activities at the transcriptional level in *E. coli*.**

Promoters were cloned to drive the sfGFP expression. Real-time quantitative PCR (qRT-PCR) was performed to determine the relative *sfGFP* transcript level in *E. coli*. All the values normalized against the internal control 16S ribosomal RNA (*rrsA*) gene. Data analyses conducted using the $2^{-\Delta\Delta Ct}$ method (Livak et al., 2001). Error bars represent standard error (SE). Data represent three biological replicates. Asterisks indicate statistically significant differences (**** $p < 0.0001$, independent samples *t*-test).

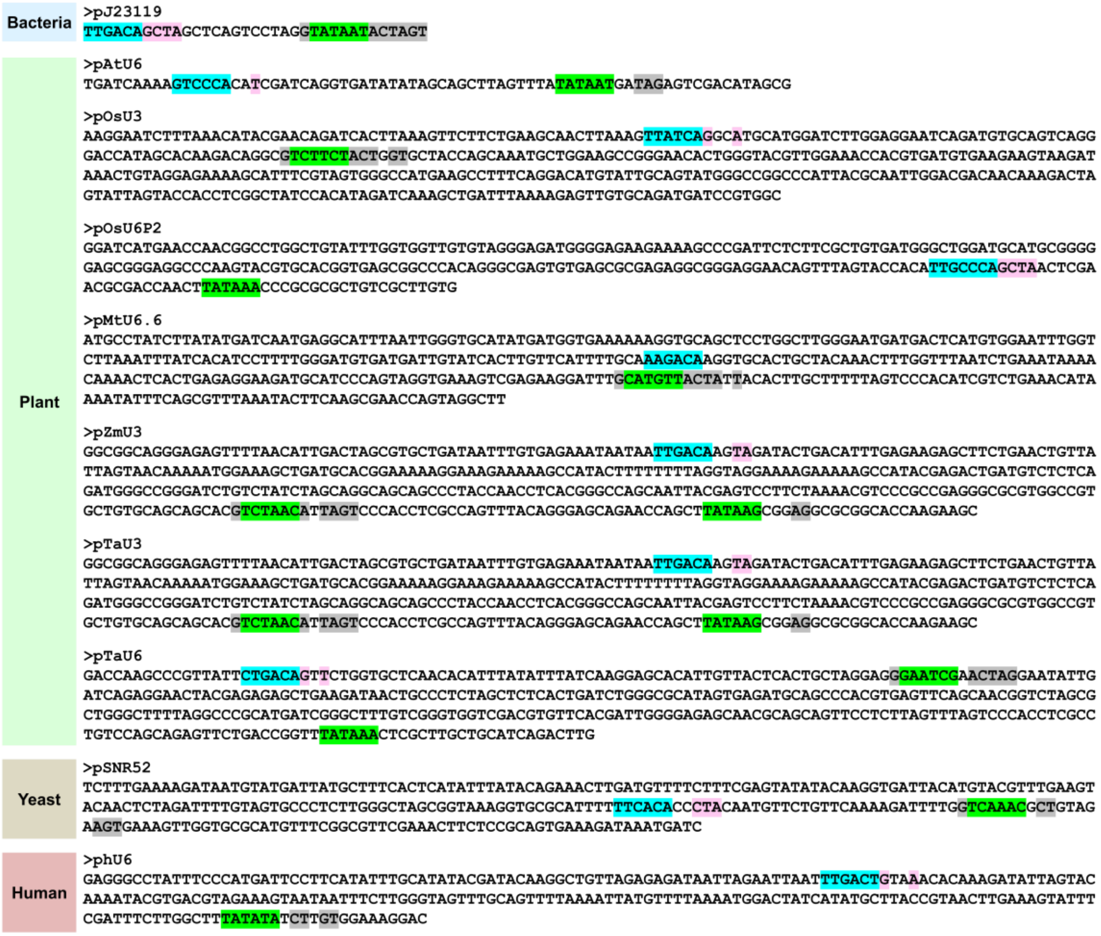


**Supplementary Figure 3. Analysis of promoters used for sgRNA expression in different organisms.**

A total of ten promoters were examined for the presence of prokaryotic promoter features, such as for conserved -10 (TATAAT) and -35 (TTGACA) elements. Seven sgRNA-expression plant-promoters include pAtU6 (Nekrasov et al., 2013), pOsU3 (Xing et al., 2014), pOsU6P2 (Lowder et al., 2015), pMtU6.6 (Jacobs et al., 2015), pZmU3 (Liang et al., 2014), pTaU3 (Lin et al., 2020), and pTaU6 (Shan et al., 2013). The sgRNA expression promoters for bacteria, yeast, and human consisted of pJ23119, pSNR52 (DiCarlo et al., 2013), and phU6 (Fu et al., 2013), respectively.

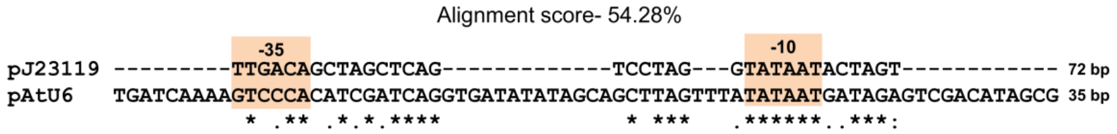


**Supplementary Figure 4. Promoter sequence alignment used for sgRNA expression in bacteria (J23119) and plant (Arabidopsis U6).**

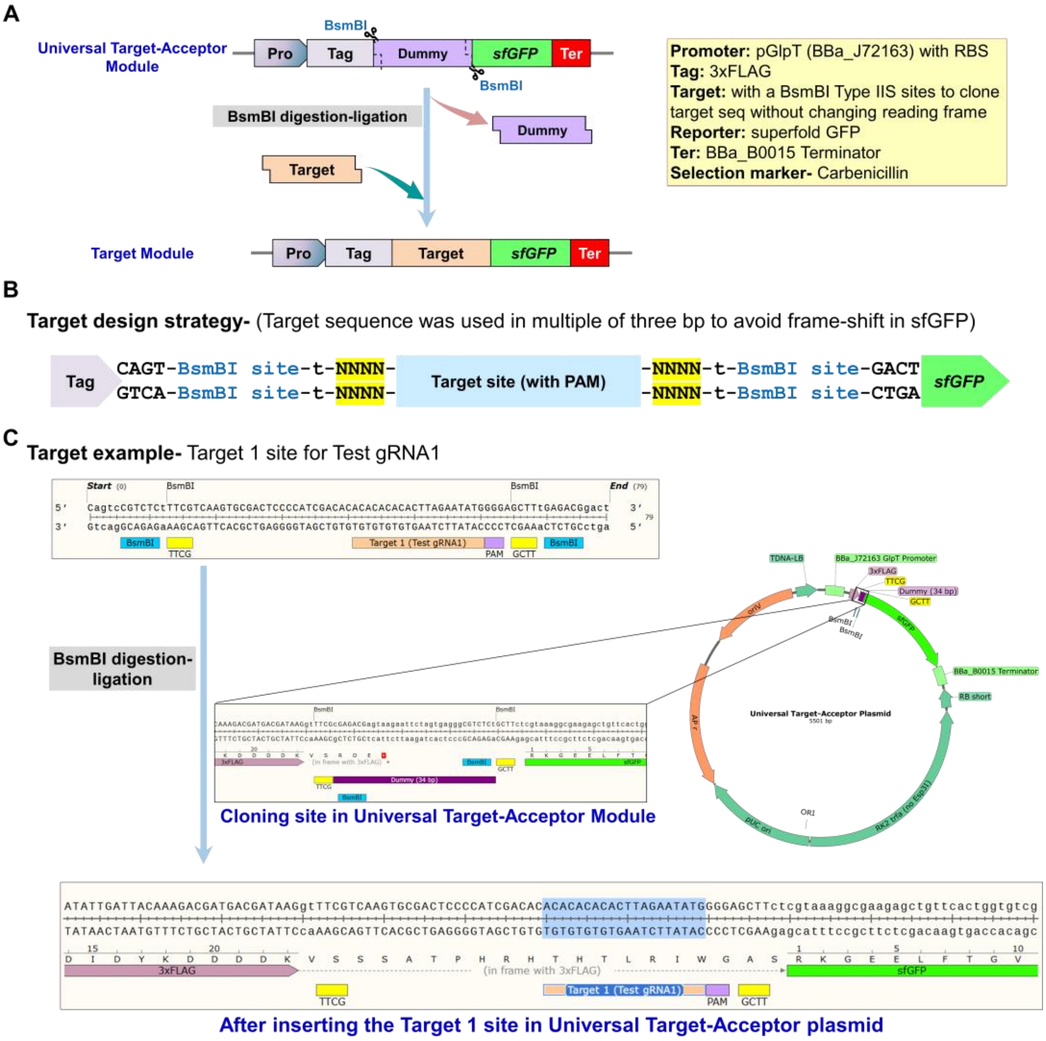


**Supplementary Figure 5. Scheme followed for cloning the target regions as DNA templates for gRNA binding and subsequently CRISPR-BE editing.**

**(A)** Schematic representation of cloning region in the universal target-acceptor module. Different components used for assembly are summarized in the yellow box. Cloning of the desired target region into a universal target-acceptor through type IIS (BsmBI) enzyme digestion-ligation permits the assessment of any intended gRNA. **(B)** The intended 23 bp target sequence (20 bp guide + 3 bp PAM) was cloned using BsmBI type IIS sites in such a manner so that codons will be in frame with the *sfGFP* sequence. The *sfGFP* sequence facilitates the screening of successfully cloned plasmids depending on the presence or absence of fluorescence. **c.** Example of target cloning for Test gRNA1 is shown.

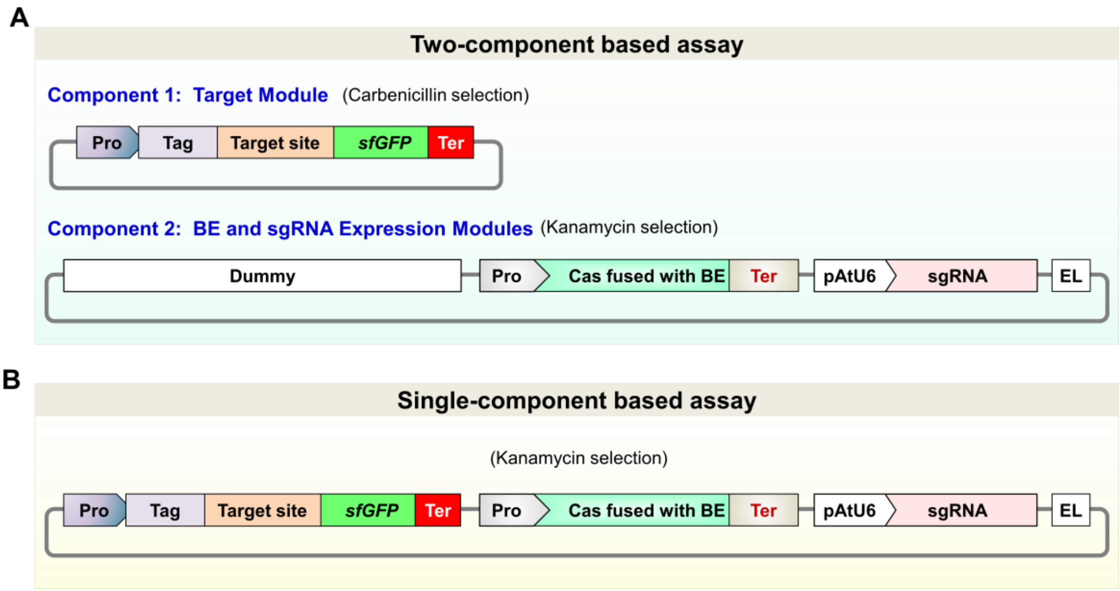


**Supplementary Figure 6. Approaches analyzed for the establishment of base editing-based IRI-CCE platform.**

**(A)** Two-component-based assay comprises two independent plasmids. The first component consisted of the target region-expression plasmid. The second component consists of an assemblage of Promoter-sgRNA and Promoter-nCas9-PmCDA1-Linker-1xUGI-Terminator (nCas9-PmCDA1) in a plasmid vector. **(B)** A single-component assay system composed of all three parts assembled into a single plasmid, including target region, Promoter-sgRNA, and nCas9-PmCDA1.

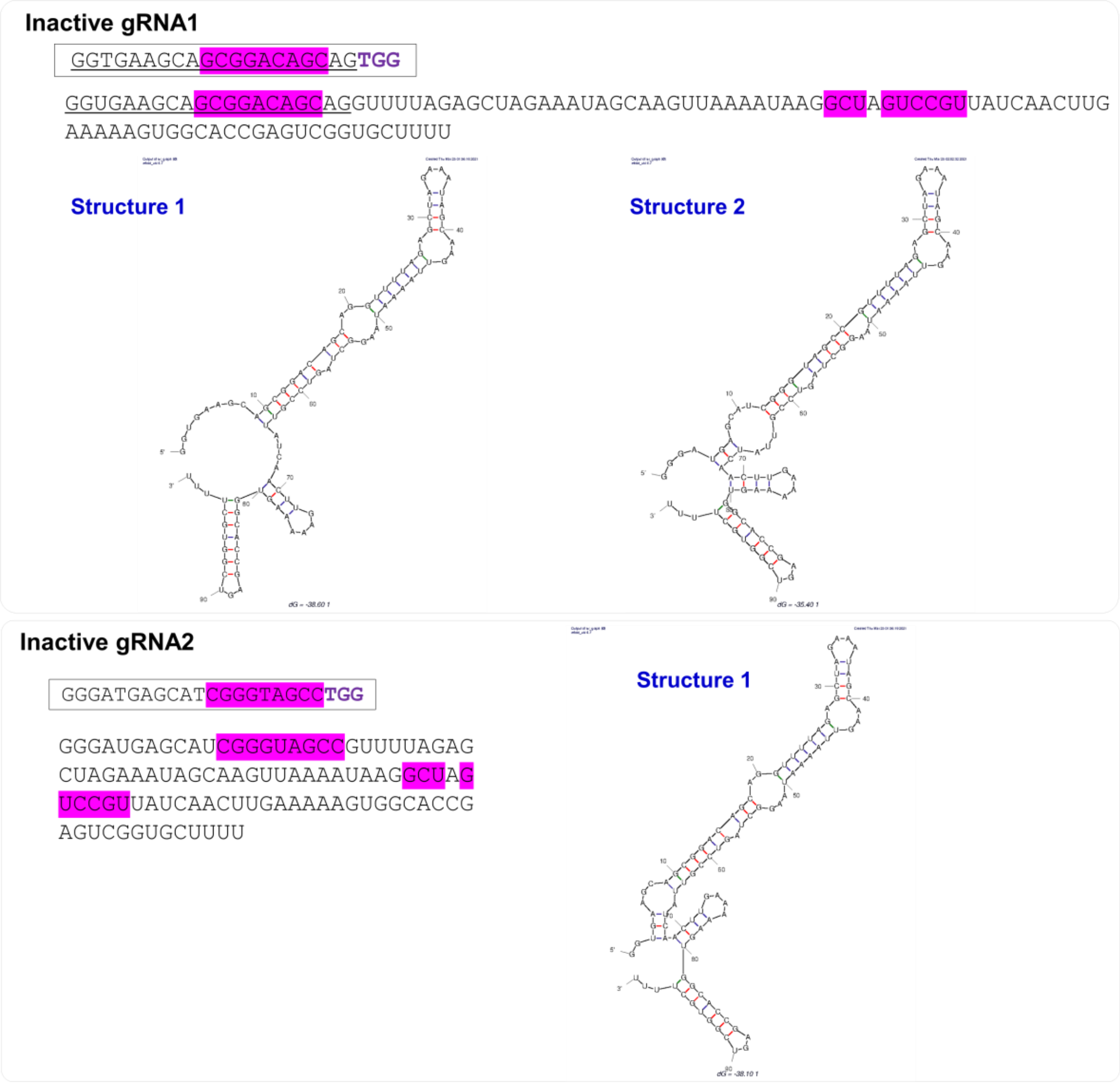


**Supplementary Figure 7. Features of inactive gRNAs from Figure 5.**

Both the gRNAs form stable interactions with the scaffold region that allow binding with Cas9, but it prevents target DNA recognition by the Cas9-sgRNA complex (Thyme et al., 2016). Secondary structures were predicted using the mfold v2.3 server using default parameters (Zuker 2003).

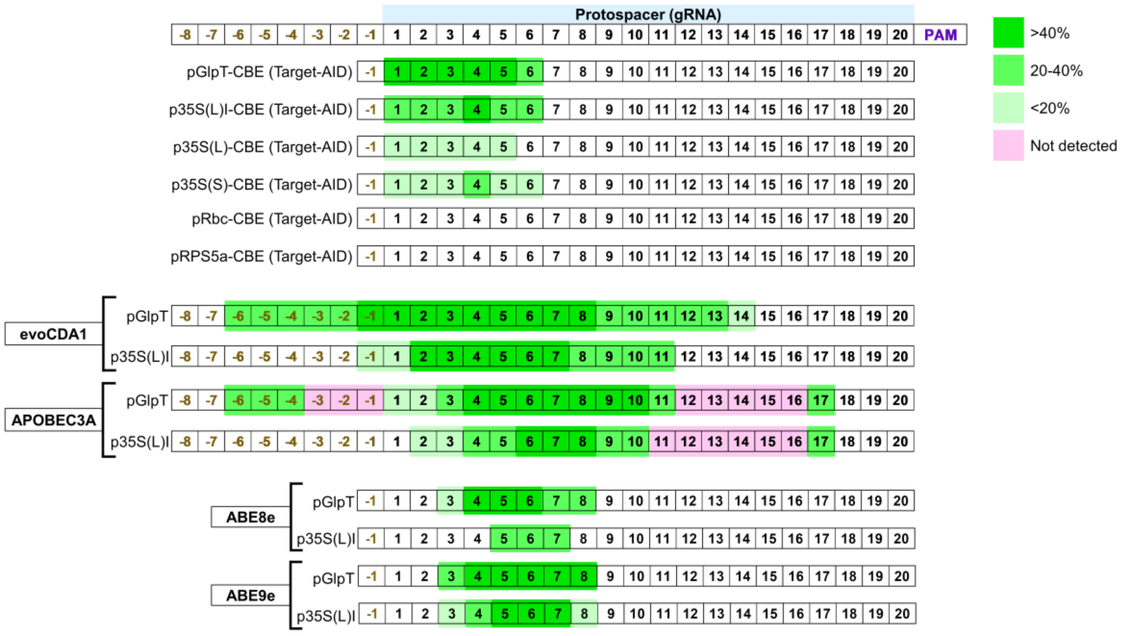


**Supplementary Figure 8. Editing windows for CBEs (PmCDA1, evoCDA1, APOBEC3A) and ABE reported in the present study expressed under the promoters of different strengths.**

**Supplementary Table 1.** *Escherichia coli* strains used in the present study.

| Strain | Genotype |
|---|---|
| 10-beta | Δ(ara-leu) 7697 araD139 fhuA ΔlacX74 galK16 galE15 e14- ϕ80dlacZΔM15 recA1 relA1 endA1 nupG rpsL (StrR) rph spoT1 Δ(mrr-hsdRMS-mcrBC) |
| DH5α | F– φ80lacZΔ M15 Δ (lacZYA-argF) U169 recA1 endA1 hsdR17 (rK– mK+) phoA supE44 λ- thi–1 gyrA96 relA1 |
| BL21(DE3) | fhuA2 [lon] ompT gal (λ DE3) [dcm] ΔhsdS λ DE3=λ sBamHIo ΔEcoRI-B int::(lacI::PlacUV5::T7 gene1) i21 Δnin5 |
| DB3.1 | gyrA462 endA1 Δ(sr1-recA) mcrB mrr hsdS20 glnV44 (=supE44) ara14 galK2 lacY1 proA2 rpsL20 xyl5 leuB6 mtl1 |

**Supplementary Table 2.** Primer sequences used for cloning and sequencing are summarized.

| **Name** | **Sequence (5'-3')** | **Description** |
|---|---|---|
| **gRNA oligos** | | |
| Test gRNA1-F | TGTGGTCTCAATTGACACACACACTTAGAATCTGGTTTTAGAGCTAGAAATAGCAAGTTAAAAT | Test gRNA1 PCR and cloning |
| Test gRNA2-F | TGTGGTCTCAATTGCACACACACATTAGAATCTGGTTTTAGAGCTAGAAATAGCAAGTTAAAAT | Test gRNA2 PCR and cloning |
| Test gRNA3-F | TGTGGTCTCAATTGAAACCCCTCCCCCACCAGCGTTTTAGAGCTAGAAATAGCAAGTTAAAAT | Test gRNA3 PCR and cloning |
| GL1-gRNA1-F | TGTGGTCTCAATTGGGAAAAGTTGTAGACTGAGAGTTTTAGAGCTAGAAATAGCAAGTTAAAAT | GL1-gRNA1 PCR and cloning |
| SlMlo1-gRNA2-F | TGTGGTCTCAATTGTACAAAGTTAATCAAGAATGTTTTAGAGCTAGAAATAGCAAGTTAAAAT | SlMlo1-gRNA2 PCR and cloning |
| SlPelo-gRNA1-F | TGTGGTCTCAATTGTCCAGCATCATTCAGTTGTGGTTTTAGAGCTAGAAATAGCAAGTTAAAAT | SlPelo-gRNA1 PCR and cloning |
| Inactive gRNA1-F | TGTGGTCTCAATTGGGTGAAGCAGCGGACAGCAGGTTTTAGAGCTAGAAATAGCAAGTTAAAAT | Inactive gRNA1 PCR and cloning |
| Inactive gRNA2-F | TGTGGTCTCAATTGGGGATGAGCATCGGGTAGCCGTTTTAGAGCTAGAAATAGCAAGTTAAAAT | Os- gRNA22 PCR and cloning |
| Os- gRNA22-F | TGTGGTCTCAATTGCTTAATTTCTGTTAGGTTCTGTTTTAGAGCTAGAAATAGCAAGTTAAAAT | Os- gRNA23 PCR and cloning |
| Os- gRNA23-F | TGTGGTCTCAATTGAGTACAAGTTTATTTTTTTCGTTTTAGAGCTAGAAATAGCAAGTTAAAAT | *galK* gRNA1 PCR and cloning |
| *galK* gRNA1-F | TGTGGTCTCAATTGCAACTGCGTAACAACAGCTTGTTTTAGAGCTAGAAATAGCAAGTTAAAAT | *rpoB* gRNA1 PCR and cloning |
| *rpoB* gRNA1-F | TGTGGTCTCAATTGGGTCCATAAACTGAGACAGCGTTTTAGAGCTAGAAATAGCAAGTTAAAAT | *rppH* gRNA1 PCR and cloning |
| *rppH* gRNA1-F | TGTGGTCTCAATTGATCGCCAGGGGCAGGTAATGGTTTTAGAGCTAGAAATAGCAAGTTAAAAT | *rppH* gRNA2 PCR and cloning |
| *rppH* gRNA2-F | TGTGGTCTCAATTGAGGGGCAGGTAATGTGGGCCGTTTTAGAGCTAGAAATAGCAAGTTAAAAT | *rppH* gRNA3 PCR and cloning |
| *rppH* gRNA3-F | TGTGGTCTCAATTGTCCTGGCAATTTCCGCAAGGGTTTTAGAGCTAGAAATAGCAAGTTAAAAT | PCR and cloning |
| gRNA Rev | TGTGGTCTCAAGCGTAATGCCAACTTTGTAC | gRNA PCR and cloning |
| **Target oligos** | | |
| Target (Test gRNA1)-F | CAGTCCGTCTCTTTCGTCAAGTGCGACTCCCCATCGACACACACACACACTTAGAATATGGGGAGCTTTGAGACGGACT | Target (Test gRNA1) |
| Target (Test gRNA1)-R | AGTCCGTCTCAAAGCTCCCCATATTCTAAGTGTGTGTGTGTGTCGATGGGGAGTCGCACTTGACGAAAGAGACGGACTG | Target (Test gRNA1) |
| Target (Test gRNA2)-F | CAGTCCGTCTCTTTCGTCAAGTGCGACTCCCCATCGACACCACACACACATTAGAATCTGGGGAGCTTTGAGACGGACT | Target (Test gRNA2) |

| | | |
|---|---|---|
| Target (Test gRNA2)-R | AGTCCGTCTCAAAGCTCCCCAGATTCTAATGTGTGTGTGGTGTCGA TGGGGAGTCGCACTTGACGAAAGAGACGGACTG | Target (Test gRNA2) |
| Target (Test gRNA3)-F | CAGTCCGTCTCTTTCGTCAAGTGCGACTCCCCATCGACACAAACCC CTCCCCCCACCAGCCGGAGCTTTGAGACGGACT | Target (Test gRNA3) |
| Target (Test gRNA3)-R | AGTCCGTCTCAAAGCTCCGGCTGGTGGGGGGAGGGGTTTGTGTCG ATGGGGAGTCGCACTTGACGAAAGAGACGGACTG | Target (Test gRNA3) |
| Target (AtGL1-gRNA1)-F | CAGTCGAAGACAATAAGGGAAAAGTTGTAGACTGAGATGGAAGT GTTGTCTTCGACTG | Target (AtGL1-gRNA1) |
| Target (AtGL1-gRNA1)-R | CAGTCGAAGACAATAAGGATCAAAGACCATCTGAATTTGGAAGTG TTGTCTTCGACTG | Target (AtGL1-gRNA1) |
| Target (SlMlo1-gRNA2)-F | CAGTCCGTCTCTTTCGGAATTGTACAAAGTTAATCAAGAATAGGG GGCTTTGAGACGGACT | Target (SlMlo1-gRNA2) |
| Target (SlMlo1-gRNA2)-R | AGTCCGTCTCAAAGCCCCCTATTCTTGATTAACTTTGTACAATTCC GAAAGAGACGGACTG | Target (SlMlo1-gRNA2) |
| Target (SlPelo-gRNA1)-F | CAGTCCGTCTCTTTCGATATATTAGACATCCAGCATCATTCAGTTG TGGGGAGCTTTGAGACGGACT | Target (SlPelo-gRNA1) |
| Target-(SlPelo-gRNA1)-R | AGTCCGTCTCAAAGCTCCCCACAACTGAATGATGCTGGATGTCTA ATATATCGAAAGAGACGGACTG | Target (SlPelo-gRNA1) |
| Target-(Inactive gRNA1)-F | CAGTCCGTCTCTTTCGCCCCATCGACACGGTGAAGCAGCGGACAG CAGTGGAGCTTTGAGACGGACT | Target-(Inactive gRNA1) |
| Target-(Inactive gRNA1)-R | AGTCCGTCTCAAAGCTCCACTGCTGTCCGCTGCTTCACCGTGTCGA TGGGGCGAAAGAGACGGACTG | Target-(Inactive gRNA1) |
| Target-(Inactive gRNA2)-F | CAGTCCGTCTCTTTCGCCCCATCGACACGGGATGAGCATCGGGTA GCCTGGAGCTTTGAGACGGACT | Target-(Inactive gRNA2) |
| Target-(Inactive gRNA2)-R | AGTCCGTCTCAAAGCTCCAGGCTACCCGATGCTCATCCCGTGTCGA TGGGGCGAAAGAGACGGACTG | Target-(Inactive gRNA2) |
| Target-(Os- gRNA22)-F | CAGTCCGTCTCTTTCGCTAATTCATGGACTTAATTTCTGTTAGGTTC TTGGAGCTTTGAGACGGACT | Target-(Os- gRNA22) |
| Target-(Os- gRNA22)-R | AGTCCGTCTCAAAGCTCCAAGAACCTAACAGAAATTAAGTCCATG AATTAGCGAAAGAGACGGACTG | Target-(Os- gRNA22) |
| Target-(Os- gRNA23)-F | CAGTCCGTCTCTTTCGATGGCATACTGAAGTACAAGTTTATTTTTT TCTGGAGCTTTGAGACGGACT | Target-(Os- gRNA23) |
| Target-(Os- gRNA23)-R | AGTCCGTCTCAAAGCTCCAGAAAAAAATAAACTTGTACTTCAGTA TGCCATCGAAAGAGACGGACTG | Target-(Os- gRNA23) |
| **qRT-PCR** | | |
| qSfGFP-F | GGTGAAGGTGACGCAACTAA | qRT-PCR of *sfGFP* |
| qSfGFP-R | GCAAAGCACTGAACACCATAAG | qRT-PCR of *sfGFP* |
| qRrsA-F | CTCTTGCCATCGGATGTGCCCA | qRT-PCR of 16S rRNA |
| qRrsA-R | CCAGTGTGGCTGGTCATCCTCTCA | qRT-PCR of 16S rRNA |
| **Target PCR and Sanger Sequencing** | | |
| L1-F1 | GATGGGCTGCCTGTATCGAGT | Target region seq |
| galk9-F | GCCAACGCATTTGGCTACCCTG | *galK* PCR and seq |

| | | |
|---|---|---|
| galk9-R | CATGCGCAACAGCGTTGAACTC | *galK* PCR |
| rpoB-F | CCTCGGCAACCGTCGTATCC | *rpoB* PCR and seq |
| rpoB-R | CCTGGGCGATAACGTAGTTGC | *rpoB* PCR |
| rpph-F | CGGCTATCCACCCCTTCCTCTG | *rppH* PCR and seq |
| rpph-R1 | GATTTCTGCATCGCCGCTCACC | *rppH* PCR |
| **PCR** | | |
| pGlpT-F1 | GAGAAGACTTGGAGGAAAGTGAAACGTGATTTCATGCGTC | pGlpT (Pro-5U) PCR |
| pGlpT-R1 | ACTGAAGACAACATTCTAGTATTTCTCCTCTTTCTCTAG | pGlpT (Pro-5U) PCR |
| pGlpT-R2 | ACTGAAGACAAATGGCTAGTATTTCTCCTCTTTCTCTAG | pGlpT (Pro-5Uf) PCR |
| Flag-F1 | ACTGAAGACTTAATGGACTATAAGGACCACGACGGAGACTACAA GGATCATGATATTGATTACAAAGACGATGACGATAAGGTTTCGAA GTCTTCACTG | Design of universal target acceptor |
| Flag-R1 | CAGTGAAGACTTCGAAACCTTATCGTCATCGTCTTTGTAATCAATA TCATGATCCTTGTAGTCTCCGTCGTGGTCCTTATAGTCCATTAAGT CTTCAGT | Design of universal target acceptor |
| DummyCT-F1 | ACTGAAGACTTTTCGCGAGACGAGTAAGAATTCTAGTGAGGGCGT CTCTGCTTAAGTCTTCACTG | Design of universal target acceptor |
| DummyCT-R1 | CAGTGAAGACTTAAGCAGAGACGCCCTCACTAGAATTCTTACTCG TCTCGCGAAAAGTCTTCAGT | Design of universal target acceptor |
| sfGFP-F1 | GAGAAGACTTGCTTCTCGTAAAGGCGAAGAGCTGTTCACTGGT | Design of universal target acceptor |
| sfGFP-R1 | ACTGAAGACAAAGCGTATAAACGCAGAAAGGCCCACCCGAA | Design of universal target acceptor |
| L3S2P2-F1 | GAGAAGACTTGCTTCTCGGTACCAAATTCCAGAAAAGAGGCCTCC CGAAAGGGGGGCCTTTTTCGTTTTGGTCCCGCTAAGTCTTCACT | TerL3S2P21 (3U-Ter) cloning |
| L3S2P2-R1 | AGTGAAGACTTAGCGGGACCAAAACGAAAAAGGCCCCCCTTTCG GGAGGCCTCTTTTCTGGAATTTGGTACCGAGAAGCAAGTCTTCTC | TerL3S2P21 (3U-Ter) cloning |
| eCDA-F1 | GCATGAAGACTTCCATGAAACGGACAGCCGACGGAAGCGAGTTC GAGTCA | evoCDA domestication (NT1) |
| eCDA-R1 | CAGTGAAGACTTGCTCTCTTCAGTGTTTTCTCCAGCCACCGATTC | evoCDA domestication (NT1) |
| eCDA-F2 | GCATGAAGACTTGAGCCGAGAAGCGGAGAAGCGA | evoCDA domestication (NT1) |
| eCDA-R2 | CAGTGAAGACTTCATTCCGCTGCCGCCGCTGCTGCCGCCACT | evoCDA domestication (NT1) |
| 2xU-F1 | GCATGAAGACTTGCTTCTAGCGGGGGGAGCACTAATCTGAGCGAC A | 2xUGI cloning |
| 2xU-R1 | CAGTGAAGACTTTACCTTAGACTTTCCTCTTCTTCTTGGGCTCGA | 2xUGI cloning |
| A3A-F1 | GCATGAAGACTTCCATGAAACGGACAGCCGACGGAAGCGAGTTC GAGTCACCAAAGAAGAAGCGGAAAGTCGAGGCCAGCCCGGCTAG CGGCCCAAGG | APOBEC3A cloning (NT1) |

| | | |
|---|---|---|
| A3A-R1 | CAGTGAAGACTTCATTCCCTTAAGAGATTCTGGGGTGGCCGAC | APOBEC3A cloning (NT1) |

**Supplementary Table 3.** Plasmids used in the present work.

| Name | Feature | Source |
|---|---|---|
| pICH86966::AtU6p::sgRNA_PDS | PCR template for sgRNA | Nekrasov et al., 2013 |
| pYTK001 (Addgene #65108) | PCR template for pGlpT, sfGFP, and sfGFP-Ter | Lee et al., 2015 |
| pICSL01009::AtU6p | Source of AtU6 promoter | Nekrasov et al., 2013 |
| pEc1-sfGFP-TerL3S2P21 | L1 module for promoter activity analysis | This work |
| pGlpT-sfGFP-TerL3S2P21 | L1 module for promoter activity analysis | This work |
| p35S(L)I-sfGFP-Ter35S | L1 module for promoter activity analysis | This work |
| p35S(L)-sfGFP-Ter35S | L1 module for promoter activity analysis | This work |
| p35S(S)-sfGFP-Ter35S | L1 module for promoter activity analysis | This work |
| pRbc-sfGFP-Ter35S | L1 module for promoter activity analysis | This work |
| pRPS5a-sfGFP-TerHSP | L1 module for promoter activity analysis | This work |
| pICH51266 (Addgene #50267) | Source of p35S(L) | Engler et al., 2014 |
| pICH51277 (Addgene #50268) | Source of p35S(S) | Engler et al., 2014 |
| pICH45195 (Addgene #50275) | Source of pRbc | Engler et al., 2014 |
| pICH41414 (Addgene #50337) | Source of Ter35S | Engler et al., 2014 |
| pKI1.1R (Addgene #85808) | Source of pRPS5a and TerHSP | Tsutsui and Higashiyama, 2017 |
| Level 1 hCas9 module (Addgene #49771) | Source of Cas9 | Nekrasov et al., 2013 |
| pGlpT-nCas9-PmCDA1-1xUGI-TerL3S2P21 | L1 module for Target-AID-based C-to-T editing | This work |
| p35S(L)I-nCas9-PmCDA1-1xUGI-Ter35S | L1 module for Target-AID-based C-to-T editing | This work |
| p35S(S)-nCas9-PmCDA1-1xUGI-Ter35S | L1 module for Target-AID-based C-to-T editing | This work |
| p35S(L)-nCas9-PmCDA1-1xUGI-Ter35S | L1 module for Target-AID-based C-to-T editing | This work |
| pRbc-nCas9-PmCDA1-1xUGI-Ter35S | L1 module for Target-AID-based C-to-T editing | This work |
| pRPS5a-sfGFP-PmCDA1-1xUGI-TerHSP | L1 module for Target-AID-based C-to-T editing | This work |
| PmCDA1-1xUGI (Addgene #79620) | Source of PmCDA1-1xUGI | Nishida et al., 2016 |
| evoCDA1 pBT277 (Addgene #122608) | Source of evoCDA1 and 2xUGI | Thuronyi et al., 2019 |
| A3A-PBE-ΔUGI (Addgene #119770) | Source of APOBEC3A | Zong et al., 2018 |

| | | |
|---|---|---|
| pGlpT-evoCDA1-nCas9-2xUGI-TerL3S2P21 | L1 module for evoCDA1-based C-to-T editing | This work |
| p35S(L)I-evoCDA1-nCas9-2xUGI-Ter35S | L1 module for evoCDA1-based C-to-T editing | This work |
| pGlpT-A3A-nCas9-2xUGI-TerL3S2P21 | L1 module for APOBEC3A-based C-to-T editing | This work |
| p35S(L)I-A3A-nCas9-2xUGI-Ter35S | L1 module for APOBEC3A-based C-to-T editing | This work |
| pGlpT-ABE8e-nCas9-TerL3S2P21 | L1 module for ABE8e-based A-to-G editing | This work |
| p35S(L)I-ABE8e-nCas9-Ter35S | L1 module for ABE8e-based A-to-G editing | This work |

**Supplementary Table 4.** DNA sequences of CRISPR and BE components

**>nCas9**

ATGGATAAAAAGTATTCTATTGGTTTAGCCATCGGCACTAATTCCGTTGGATGGGCTGTCATAACCGATGAATACAAA
GTACCTTCAAAGAAATTTAAGGTGTTGGGGAACACAGACCGTCATTCGATTAAAAAGAATCTTATCGGTGCCCTCCT
ATTCGATAGTGGCGAAACGGCAGAGGCGACTCGCCTGAAACGAACCGCTCGGAGAAGGTATACACGTCGCAAGAAC
CGAATATGTTACTTACAAGAAATTTTTAGCAATGAGATGGCCAAAGTTGACGATTCTTTCTTTCACCGTTTGGAAGAG
TCCTTCCTTGTCGAAGAGGACAAGAAACATGAACGGCACCCCATCTTTGGAAACATAGTAGATGAGGTGGCATATCA
TGAAAAGTACCCAACGATTTATCACCTCAGAAAAAAGCTAGTTGACTCAACTGATAAAGCGGACCTGAGGTTAATCT
ACTTGGCTCTTGCCCATATGATAAAGTTCCGTGGGCACTTTCTCATTGAGGGTGATCTAAATCCGGACAACTCGGATG
TCGACAAACTGTTCATCCAGTTAGTACAAACCTATAATCAGTTGTTTGAAGAGAACCCTATAAATGCAAGTGGCGTG
GATGCGAAGGCTATTCTTAGCGCCCGCCTCTCTAAATCCCGACGGCTAGAAAACCTGATCGCACAATTACCCGGAGA
GAAGAAAAATGGGTTGTTCGGTAACCTTATAGCGCTCTCACTAGGCCTGACACCAAATTTTAAGTCGAACTTCGACTT
AGCTGAAGATGCCAAATTGCAGCTTAGTAAGGACACGTACGATGACGATCTCGACAATCTACTGGCACAAATTGGAG
ATCAGTATGCGGACTTATTTTTGGCTGCCAAAAACCTTAGCGATGCAATCCTCCTATCTGACATACTGAGAGTTAATA
CTGAGATTACCAAGGCGCCGTTATCCGCTTCAATGATCAAAAGGTACGATGAACATCACCAAGACTTGACACTTCTC
AAGGCCCTAGTCCGTCAGCAACTGCCTGAGAAATATAAGGAAATATTCTTTGATCAGTCGAAAAACGGGTACGCAGG
TTATATTGACGGCGGAGCGAGTCAAGAGGAATTCTACAAGTTTATCAAACCCATATTAGAGAAGATGGATGGGACGG
AAGAGTTGCTTGTAAAACTCAATCGCGAAGATCTACTGCGAAAGCAGCGGACTTTCGACAACGGTAGCATTCCACAT
CAAATCCACTTAGGCGAATTGCATGCTATACTTAGAAGGCAGGAGGATTTTTATCCGTTCCTCAAAGACAATCGTGA
AAAGATTGAGAAAATCCTAACCTTTCGCATACCTTACTATGTGGGACCCCTGGCCCGAGGGAACTCTCGGTTCGCATG
GATGACAAGAAAGTCCGAAGAAACGATTACTCCATGGAATTTTGAGGAAGTTGTCGATAAAGGTGCGTCAGCTCAAT
CGTTCATCGAGAGGATGACCAACTTTGACAAGAATTTACCGAACGAAAAAGTATTGCCTAAGCACAGTTTACTTTAC
GAGTATTTCACAGTGTACAATGAACTCACGAAAGTTAAGTATGTCACTGAGGGCATGCGTAAACCCGCCTTTCTAAG
CGGAGAACAGAAGAAAGCAATAGTAGATCTGTTATTCAAGACCAACCGCAAAGTGACAGTTAAGCAATTGAAAGAG
GACTACTTTAAGAAAATTGAATGCTTCGATTCTGTCGAGATCTCCGGGGTAGAAGATCGATTTAATGCGTCACTTGGT
ACGTATCATGACCTCCTAAAGATAATTAAAGATAAGGACTTCCTGGATAACGAAGAGAATGAAGATATCTTAGAAGA
TATAGTGTTGACTCTTACCCTCTTTGAAGATCGGGAAATGATTGAGGAAAGACTAAAAACATACGCTCACCTGTTCGA
CGATAAGGTTATGAAACAGTTAAAGAGGCGTCGCTATACGGGCTGGGGACGATTGTCGCGGAAACTTATCAACGGG
ATAAGAGACAAGCAAAGTGGTAAAACTATTCTCGATTTTCTAAAGAGCGACGGCTTCGCCAATAGGAACTTTATGCA
GCTGATCCATGATGACTCTTTAACCTTCAAAGAGGATATACAAAAGGCACAGGTTTCCGGACAAGGGGACTCATTGC
ACGAACATATTGCGAATCTTGCTGGTTCGCCAGCCATCAAAAAGGGCATACTCCAGACAGTCAAAGTAGTGGATGAG
CTAGTTAAGGTCATGGGACGTCACAAACCGGAAAACATTGTAATCGAGATGGCACGCGAAAATCAAACGACTCAGA
AGGGGCAAAAAAACAGTCGAGAGCGGATGAAGAGAATAGAAGAGGGTATTAAAGAACTGGGCAGCCAGATCTTAA
AGGAGCATCCTGTGGAAAATACCCAATTGCAGAACGAGAAACTTTACCTCTATTACCTACAAAATGGAAGGGACATG
TATGTTGATCAGGAACTGGACATAAACCGTTTATCTGATTACGACGTCGATCACATTGTACCCCAATCCTTTTTGAAG
GACGATTCAATCGACAATAAAGTGCTTACACGCTCGGATAAGAACCGAGGGAAAAGTGACAATGTTCCAAGCGAGG
AAGTCGTAAAGAAAATGAAGAACTATTGGCGGCAGCTCCTAAATGCGAAACTGATAACGCAAAGAAAGTTCGATAA
CTTAACTAAAGCTGAGAGGGGTGGCTTGTCTGAACTTGACAAGGCCGGATTTATTAAACGTCAGCTCGTGGAAACCC
GCCAAATCACAAAGCATGTTGCACAGATACTAGATTCCCGAATGAATACGAAATACGACGAGAACGATAAGCTGATT
CGGGAAGTCAAAGTAATCACTTTAAAGTCAAAATTGGTGTCGGACTTCAGAAAGGATTTTCAATTCTATAAAGTTAG
GGAGATAAATAACTACCACCATGCGCACGACGCTTATCTTAATGCCGTCGTAGGGACCGCACTCATTAAGAAATACC
CGAAGCTAGAAAGTGAGTTTGTGTATGGTGATTACAAAGTTTATGACGTCCGTAAGATGATCGCGAAAAGCGAACAG
GAGATAGGCAAGGCTACAGCCAAATACTTCTTTTATTCTAACATTATGAATTTCTTTAAGACGGAAATCACTCTGGCA
AACGGAGAGATACGCAAACGACCTTTAATTGAAACCAATGGGGAGACAGGTGAAATCGTATGGGATAAGGGCCGGG
ACTTCGCGACGGTGAGAAAAGTTTTGTCCATGCCCCAAGTCAACATAGTAAAGAAAACTGAGGTGCAGACCGGAGG
GTTTTCAAAGGAATCGATTCTTCCAAAAAGGAATAGTGATAAGCTCATCGCTCGTAAAAAGGACTGGGACCCGAAAA
AGTACGGTGGCTTCGATAGCCCTACAGTTGCCTATTCTGTCCTAGTAGTGGCAAAAGTTGAGAAGGGAAAATCCAAG
AAACTGAAGTCAGTCAAAGAATTATTGGGGATAACGATTATGGAGCGCTCGTCTTTTGAAAAGAACCCCATCGACTT
CCTTGAGGCGAAAGGTTACAAGGAAGTAAAAAAGGATCTCATAATTAAACTACCAAAGTATAGTCTGTTTGAGTTAG
AAAATGGCCGAAAACGGATGTTGGCTAGCGCCGGAGAGCTTCAAAAGGGGAACGAACTCGCACTACCGTCTAAATA
CGTGAATTTCCTGTATTTAGCGTCCCATTACGAGAAGTTGAAAGGTTCACCTGAAGATAACGAACAGAAGCAACTTTT
TGTTGAGCAGCACAAACATTATCTCGACGAAATCATAGAGCAAATTTCGGAATTCAGTAAGAGAGTCATCCTAGCTG
ATGCCAATCTGGACAAAGTATTAAGCGCATACAACAAGCACAGGGATAAACCCATACGTGAGCAGGCGGAAAATAT
TATCCATTTGTTTACTCTTACCAACCTCGGCGCTCCAGCCGCATTCAAGTATTTTGACACAACGATAGATCGCAAACG
ATACACTTCTACCAAGGAGGTGCTAGACGCGACACTGATTCACCAATCCATCACGGGATTATATGAAACTCGGATAG
ATTTGTCACAGCTTGGGGGTGAC

**>PmCDA1-1xUGI (SH3 Linker-3xFLAG-PmCDA1-1xUGI)**

GGTGGAGGAGGTACCGGCGGTGGAGGCTCAGCAGAATACGTACGAGCTCTGTTTGACTTCAATGGGAATGACGAGG
AGGATCTCCCCTTTAAGAAGGGCGATATTCTCCGCATCAGAGATAAGCCCGAAGAACAATGGTGGAATGCCGAGGAT
AGCGAAGGGAAAAGGGGCATGATTCTGGTGCCATATGTGGAGAAATATTCCGGTGACTACAAAGACCATGATGGGG
ATTACAAAGACCACGACATCGACTACAAAGACGACGACGATAAATCAGGGATGACAGACGCCGAGTACGTGCGCAT
TCATGAGAAACTGGATATTTACACCTTCAAGAAGCAGTTCTTCAACAACAAGAAATCTGTGTCACACCGCTGCTACGT
GCTGTTTGAGTTGAAGCGAAGGGGCGAAAGAAGGGCTTGCTTTTGGGGCTATGCCGTCAACAAGCCCCAAAGTGGCA
CCGAGAGAGGAATACACGCTGAGATATTCAGTATCCGAAAGGTGGAAGAGTATCTTCGGGATAATCCTGGGCAGTTT
ACGATCAACTGGTATTCCAGCTGGAGTCCTTGCGCTGATTGTGCCGAGAAAATTCTGGAATGGTATAATCAGGAACTT
CGGGGAAACGGGCACACATTGAAAATCTGGGCCTGCAAGCTGTACTACGAGAAGAATGCCCGGAACCAGATAGGAC
TCTGGAATCTGAGGGACAATGGTGTAGGCCTGAACGTGATGGTTTCCGAGCACTATCAGTGTTGTCGGAAGATTTTCA
TCCAAAGCTCTCATAACCAGCTCAATGAAAACCGCTGGTTGGAGAAAACACTGAAACGTGCGGAGAAGTGGAGATC
CGAGCTGAGCATCATGATCCAGGTCAAGATTCTGCATACCACTAAGTCTCCAGCCGTTGGTCCCAAGAAGAAAAGAA
AAGTCGGTACCATGACCAACCTTTCCGACATCATAGAGAAGGAAACAGGCAAACAGTTGGTCATCCAAGAGTCGATA
CTCATGCTTCCTGAAGAAGTTGAGGAGGTCATTGGGAATAAGCCGGAAAGTGACATTCTCGTACACACTGCGTATGA
TGAGAGCACCGATGAGAACGTGATGCTGCTCACGTCAGATGCCCCAGAGTACAAACCCTGGGCTCTGGTGATTCAGG
ACTCTAATGGAGAGAACAAGATCAAGATGCTATAA

**>evoCDA1-XTEN**

AGTACCGACGCCGAGTACGTGCGGATCCACGAGAAGCTGGATATCTATACATTCAAGAAGCAGTTTAGCAACAATAA
GAAGTCCGTGTCTCACAGATGCTACGTGCTGTTCGAGCTGAAGCGGAGAGGAGAGAGGCGCGCCTGTTTTTGGGGCT
ATGCCGTGAACAAGCCACAGTCTGGAACCGAGAGGGGAATCCACGCAGAGATCTTCAGCATCAGGAAGGTGGAGGA
GTACCTGCGCGACAACCCCGGCCAGTTTACAATCAATTGGTATAGCTCCTGGAGCCCTTGCGCCGATTGTGCCGAGA
AGATCCTGGAGTGGTACAACCAGGAGCTGAGGGGCAATGGCCACACCCTGAAGATCTGGGTGTGCAAGCTGTACTAT
GAGAAGAACGCCAGGAATCAGATCGGCCTGTGGAACCTGCGCGACAATGGCGTGGGCCTGAACGTGATGGTGTCCG
AGCACTATCAGTGCTGTCGCAAGATCTTTATCCAGTCTAGCCACAATCAGCTGAACGAGAATCGGTGGCTGGAGAAA
ACACTGAAGAGAGCCGAGAAGCGGAGAAGCGAGCTGTCCATCATGTTTCAGGTGAAGATCCTGCACACCACAAAGT
CTCCCGCCGTGTCTGGCGGATCTAGCGGAGGATCCTCTGGCAGCGAGACACCAGGAACAAGCGAGTCAGCAACACC
AGAGAGCAGTGGCGGCAGCAGCGGCGGCAGC

**>Link-2xUGI**

TCTGGTGGTTCTGGTTCGAGCGGAGGATCCGGAGGATCTGGAGGCAGCGCTTCTAGCGGGGGGAGCACTAATCTGAG
CGACATCATTGAGAAGGAGACTGGGAAACAGCTGGTCATTCAGGAGTCCATCCTGATGCTGCCTGAGGAGGTGGAG
GAAGTGATCGGCAACAAGCCAGAGTCTGACATCCTGGTGCACACCGCCTACGACGAGTCCACAGATGAGAATGTGAT
GCTGCTGACCTCTGACGCCCCCGAGTATAAGCCTTGGGCCCTGGTCATCCAGGATTCTAACGGCGAGAATAAGATCA
AGATGCTGAGCGGAGGATCCGGAGGATCTGGAGGCAGCACCAACCTGTCTGACATCATCGAGAAGGAGACAGGCAA
GCAGCTGGTCATCCAGGAGAGCATCCTGATGCTGCCCGAAGAAGTCGAAGAAGTGATCGGAAACAAGCCTGAGAGC
GATATCCTGGTCCATACCGCCTACGACGAGAGTACCGACGAAAATGTGATGCTGCTGACATCTGACGCCCCAGAGTA
TAAGCCCTGGGCTCTGGTCATCCAGGATTCCAACGGAGAGAACAAAATCAAAATGCTG

**>APOBEC3A-XTEN**

GAGGCCAGCCCGGCTAGCGGCCCAAGGCATCTCATGGACCCGCACATCTTCACCAGCAACTTCAACAACGGCATCGG
CAGGCACAAGACCTACTTGTGCTACGAGGTGGAGAGGCTCGACAACGGAACCTCCGTGAAGATGGACCAACACAGG
GGGTTCCTCCACAACCAAGCCAAGAACCTCCTCTGCGGCTTCTACGGCAGGCACGCCGAGTTGAGGTTCCTCGACTTG
GTGCCATCCCTCCAACTCGATCCAGCCCAAATCTACCGCGTGACCTGGTTCATCTCCTGGTCCCCATGCTTCTCCTGGG
GTTGCGCCGGCGAGGTTCGGGCTTTCCTCCAAGAAAACACCCACGTCCGCCTCCGCATTTTCGCCGCCAGGATCTATG
ATTACGACCCTCTCTACAAGGAGGCCCTCCAGATGCTGCGGGACGCCGGTGCTCAGGTGAGTATCATGACCTACGAC
GAGTTCAAGCACTGCTGGGACACCTTCGTTGACCACCAGGGCTGCCCATTCCAACCATGGGACGGTCTGGATGAACA
CAGCCAAGCCTTGTCCGGCAGGCTCCGGGCCATCCTCCAAAACCAGGGGAACTCCGGGAGCGAGACGCCAGGCACC
TCCGAGTCGGCCACCCCAGAATCTCTTAAG

**>ABE8e**

ATGAGTGAGGTGGAGTTCTCTCACGAATACTGGATGCGACATGCTCTAACGCTAGCAAAACGAGCGAGGGATGAAC
GAGAGGTTCCTGTAGGAGCAGTGTTGGTTCTGAACAACAGAGTTATTGGTGAAGGTTGGAATCGTGCTATTGGGCTT
CACGACCCAACAGCCCATGCCGAAATAATGGCGCTCAGGCAAGGAGGCTTAGTAATGCAAAACTACAGATTAATCG
ACGCGACCCTGTATGTCACGTTCGAGCCATGCGTTATGTGCGCGGGCGCGATGATTCATTCTAGAATTGGAAGGGTTG
TTTTTGGGGTGAGAAATTCTAAAAGAGGTGCGGCTGGGAGTCTTATGAATGTTCTCAATTACCCTGGTATGAACCATC
GAGTGGAAATCACGGAGGGGATTTTGGCGGACGAATGTGCAGCATTGTTATGCGATTTCTATCGTATGCCTAGGCAG
GTTTTCAACGCTCAGAAGAAGGCGCAAAGTTCAATT

**> TerL3S2P21**

TCGACTCGGTACCAAATTCCAGAAAAGAGGCCTCCCGAAAGGGGGGCCTTTTTTCGTTTTGGTCCTGTT

**> sfGFP-Stop-BBa_J72163 Ter**

ATGCGTAAAGGCGAAGAGCTGTTCACTGGTGTCGTCCCTATTCTGGTGGAACTGGATGGTGATGTCAACGGTCATAA
GTTTTCCGTGCGTGGCGAGGGTGAAGGTGACGCAACTAATGGTAAACTGACGCTGAAGTTCATCTGTACTACTGGTA
AACTGCCGGTTCCTTGGCCGACTCTGGTAACGACGCTGACTTATGGTGTTCAGTGCTTTGCTCGTTATCCGGACCATA
TGAAGCAGCATGACTTCTTCAAGTCCGCCATGCCGGAAGGCTATGTGCAGGAACGCACGATTTCCTTTAAGGATGAC
GGCACGTACAAAACGCGTGCGGAAGTGAAATTTGAAGGCGATACCCTGGTAAACCGCATTGAGCTGAAAGGCATTG
ACTTTAAAGAGGACGGCAATATCCTGGGCCATAAGCTGGAATACAATTTTAACAGCCACAATGTTTACATCACCGCC
GATAAACAAAAAAATGGCATTAAAGCGAATTTTAAAATTCGCCACAACGTGGAGGATGGCAGCGTGCAGCTGGCTG
ATCACTACCAGCAAAACACTCCAATCGGTGATGGTCCTGTTCTGCTGCCAGACAATCACTATCTGAGCACGCAAAGC
GTTCTGTCTAAAGATCCGAACGAGAAACGCGATCATATGGTTCTGCTGGAGTTCGTAACCGCAGCGGGCATCACGCA
TGGTATGGATGAACTGTACAAA**TGA**CCAGGCATCAAATAAAACGAAAGGCTCAGTCGAAAGACTGGGCCTTTCGTTT
TATCTGTTGTTTGTCGGTGAACGCTCTCTACTAGAGTCACACTGGCTCACCTTCGGGTGGGCCTTTCTGCGTTTATA
